# A new duck genome reveals conserved and convergently evolved chromosome architectures of birds and mammals

**DOI:** 10.1101/2020.11.04.368910

**Authors:** Jing Li, Jilin Zhang, Jing Liu, Yang Zhou, Cheng Cai, Luohao Xu, Xuelei Dai, Shaohong Feng, Chunxue Guo, Jinpeng Rao, Kai Wei, Erich D. Jarvis, Yu Jiang, Zhengkui Zhou, Guojie Zhang, Qi Zhou

## Abstract

**Background:** Ducks have a typical avian karyotype that consists of macro- and microchromosomes, but a pair of much less differentiated ZW sex chromosomes compared to chicken. To elucidate the evolution of chromosome architectures between duck and chicken, and between birds and mammals, we produced a nearly complete chromosomal assembly of a female Pekin duck by combining long-read sequencing and multiplatform scaffolding techniques.

**Results:** The major improvement of genome assembly and annotation quality resulted from successful resolution of lineage-specific propagated repeats that fragmented the previous Illumina-based assembly. We found that the duck topologically associated domains (TAD) are demarcated by putative binding sites of the insulator protein CTCF, housekeeping genes, or transitions of active/inactive chromatin compartments, indicating the conserved mechanisms of spatial chromosome folding with mammals. There are extensive overlaps of TAD boundaries between duck and chicken, and also between the TAD boundaries and chromosome inversion breakpoints. This suggests strong natural selection on maintaining regulatory domain integrity, or vulnerability of TAD boundaries to DNA double-strand breaks. The duck W chromosome retains 2.5-fold more genes relative to chicken. Similar to the independently evolved human Y chromosome, the duck W evolved massive dispersed palindromic structures, and a pattern of sequence divergence with the Z chromosome that reflects stepwise suppression of homologous recombination.

**Conclusions:** Our results provide novel insights into the conserved and convergently evolved chromosome features of birds and mammals, and also importantly add to the genomic resources for poultry studies.

## Background

Birds have the largest species number and some of the smallest genome sizes among terrestrial vertebrates. This has attracted extensive efforts since the era of cytogenetics into elucidating the diversity of their ‘streamlined’ genomes that give rise to the tremendous phenotypic diversity[1]. The karyotype of birds exhibits two major distinctions from that of mammals: first, it comprises about 10 pairs of large to medium sized chromosomes (macrochromosomes) and about 30 pairs of much smaller sized chromosomes (microchromosomes)[2]. During the over 100 million years (MY) of avian evolution, there were few interchromosomal rearrangements among most species[3–5] except for falcons and parrots (Falconiformes and Psittaciformes)[6–9]. Among the published karyotypes of over 800 bird species, the majority of them have a similar chromosome number around 2n=80[10]. These results indicate that the chromosome evolution of birds is dominated by intrachromosomal rearrangements. Genomic comparisons between chicken, turkey, flycatcher and zebra finch[11, 12] found that birds, similar to mammals[13, 14], have fragile genomic regions that were recurrently used for mediating intrachromosomal rearrangements, and these regions seem to be associated with high recombination rates[15] and low densities of conserved non-coding elements (CNEs)[5]. However, compared to mammals[13, 14, 16], much less is known about the interspecific diversity within avian chromosomes, particularly microchromosomes (but see[5, 12]) at the sequence level, due to the scarcity of chromosome-level bird genomes.

The other major distinction between the mammalian and avian karyotypes is their sex chromosomes. Birds have a pair of female heterogametic (male ZZ, female ZW) sex chromosomes that originated from a different pair of ancestral autosomes than the eutherian XY[17, 18]. Since their divergence about 300 MY ago, sex chromosomes of birds and mammals have undergone independent stepwise suppression of homologous recombination, and produced a punctuated pattern of pairwise sequence divergence levels between the neighboring regions termed ‘evolutionary strata’[19–21]. Despite the consequential massive gene loss, both chicken W chromosome (chrW) and eutherian chrYs have been found to preferentially retain dosagesensitive genes or genes with important regulatory functions[22]. In addition, the human chrY has evolved palindromic sequences that may facilitate gene conversions between the Y-linked gene copies[23], as an evolutionary strategy to limit the functional degeneration under the nonrecombining environment[24]. Interestingly, such palindromic structures have also been reported on sex chromosomes of New World sparrows and blackbirds[25], and more recently in a plant species, the willow[26], suggesting it is a general feature of evolving sex chromosomes. Both cytogenetic work and Illumina-based genome assemblies of tens of bird species suggested that bird sex chromosomes comprise an unexpected interspecific diversity regarding both their lengths of recombining regions (pseudoautosomal regions, PAR), and their rates of gene loss[20, 27]. For example, PARs cover over two thirds of the length of ratite (e.g., emu and ostrich) sex chromosomes[28], but are concentrated at the tips of the chicken and eutherian sex chromosomes. However, so far only the chicken chrW has been well-assembled using the laborious iterative clone-based sequencing method[22], and the majority of genomic sequencing projects tend to choose a male bird to avoid the repetitive chrW. This has hampered our broad and deep understanding of the composition and evolution of avian sex chromosomes.

The Vertebrate Genomes Project (VGP) has taken advantage of the development of long-read (PacBio or Nanopore) sequencing, linked-read (10X) and high-throughput chromatin conformation capture (Hi-C) technologies to empower rapid and accurate assembly of chromosome-level genomes including the sex chromosomes, in the absence of physical maps[29]. Further, Hi-C can uncover the three-dimensional (3D) architecture of chromosomes that is segregated in active (A) and inactive (B) chromatin compartments[30], and to a finer genomic scale, topologically associated domains (TADs) as the replication and regulatory units[31]. To elucidate the evolution of avian chromosome architectures in terms of sequence composition, genomic rearrangement and 3D chromatin structure, here we utilized a modified VGP pipeline to produce a nearly complete reference genome of a female Pekin duck (*Anas platyrhynchos,* Z2 strain) with all the cutting-edge technologies mentioned above. We corroborated our reference genome through comparisons to previously published radiation hybrid (RH)[32] and fluorescence *in situ* hybridization (FISH)[33] linkage maps. We chose duck because first, as a representative species of *Anseriformes,* it diverged from *Galliformes* about 72.5 MY ago[34], providing a deep but still trackable evolutionary distance for addressing the functional consequences of genomic rearrangements on chromatin domains. Second, the duck sex chromosomes have diverged to a degree between the highly heteromorphic sex chromosomes of chicken and homomorphic sex chromosomes of emu[20, 27]. The gradient of sex chromosome divergence levels exhibited by the three bird species together constitute a chronological order for a comprehensive understanding of the entire avian sex chromosome evolution process. Finally, besides being frequently used for basic evolutionary and developmental studies[35], the duck is another key poultry species, as well as a natural reservoir of all influenza A viruses[36]. Our new duck genome has anchored over 95% of the assembled sequences onto chromosomes, with great improvements in the non-coding regions and chrW sequences. We believe it will serve an important genomic resource for future studies into the mechanisms and application of artificial selection.

### Data Description

Pekin duck (called duck from here on) has a haploid genome size estimated to be 1.41 Gb[37, 38], and a karyotype of 9 pairs of macrochromosomes (from chr1 to chr8, chrZ/chrW) and 31 pairs of microchromosomes (chr9 to chr39)[39]. The Illumina-based genome assembly of the duck (BGI1.0) was produced over seven years ago and has 25.9% of the assembled genome assigned to chromosomes, containing 3.17% of bases as gaps[36]. To *de novo* assemble the new genome, we generated 143-X genome coverage of PacBio long reads (read N50 14.3 kb from 115 SMRT cells, **Supplementary Fig. S1**), and 142-X genome coverage of 10x linked-read data from a female individual, 56-X genome coverage of BioNano map and 82-X genome coverage of Hi-C reads from two different male individuals of the same inbred duck strain (Figure 1, **Supplementary Table S1**), and assembled the genome with a modified VGP pipeline[29]. To identify the female-specific chrW sequences, we also generated 72-X genome coverage Illumina reads from a male individual of the same duck strain to compare to the previously published female reads (SRA accession number: PRJNA636121). Our primary assembly of PacBio long reads assembles the entire genome into 1,645 gapless contigs (**Supplementary Table S2**), resulting in a 14-fold reduction of contig number (1,645 vs. 227,448) and 212-fold improvement of contig continuity measured by N50 (5.5Mb vs. 26.1Kb) compared to the BGI1.0 genome (Table 1). To scaffold the contigs, we first corrected their sequence errors with 92-X genome coverage female Illumina reads, then oriented and scaffolded them into 942 scaffolds with 10X linked-reads, BioNano optical maps and Hi-C reads (see **Methods**). As Hi-C data provides linkage but not orientation information, in our final step of chromosome anchoring, we incorporated an RH linkage map [32] and reduced the scaffold number further down to 755. We however detected 69 cases of conflicts of orientation between the RH map and the Hi-C scaffolds, manifested as inversions. By carefully examining the presence/absence of raw PacBio reads, Illumina mate-pairs, and syntenic chicken/goose sequences[40, 41] spanning the breakpoints of such inversions, the majority (54 of 69) supported the Hi-C map. And we have corrected a total of 15 orientation errors within the scaffolds (**Supplementary Fig. S2**).

**Figure 1.**
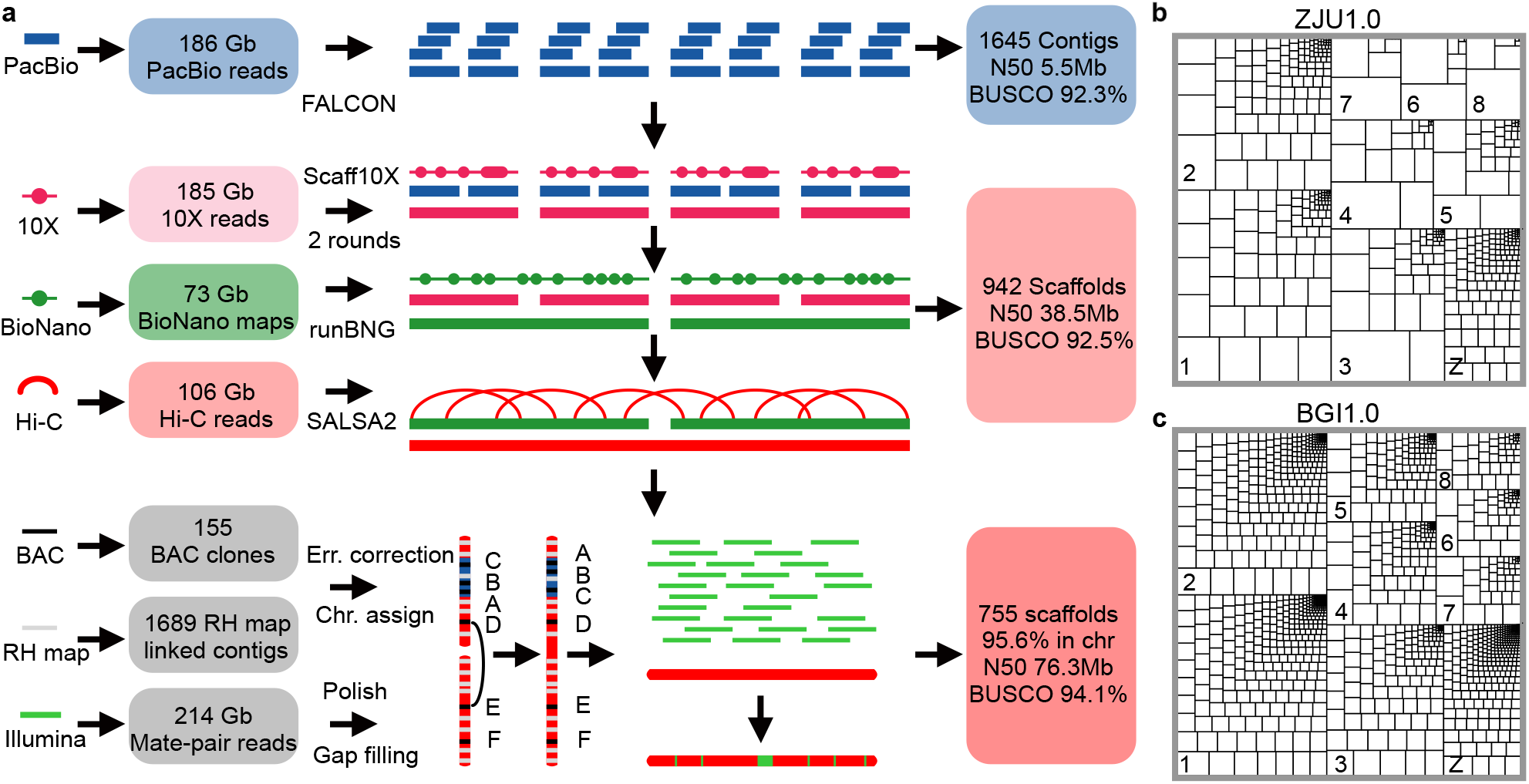
Genome assembly of a female Pekin duck. **a**. Our assembly pipeline uses high coverage PacBio long reads to generate contigs, which are then sequentially scaffolded with 10X Genomics linked reads, BioNano optical maps, Hi-C paired reads, RH maps and FISH maps, to produce a chromosome-level genome for the Pekin duck. **b, c**. Treemap comparison of contigs between ZJU1.0 and BGI1.0 versions of the duck genome. The size of each rectangle of each chromosome is scaled to that of contig sequence. The bigger and fewer the internal boxes, the more contiguous the contigs.

**Table 1.** Comparing genome assemblies of duck vs. other birds

|  | Pekin duck<br>(BGI1.0) | Pekin duck<br>(ZJU1.0) | Chicken<br>(Ncbi-6a) | Zebra finch<br>(VGP) |
| --- | --- | --- | --- | --- |
| total length (Gb) | 1.105 | 1.189 | 1.065 | 1.069 |
| #contigs | 227,448 | 1,645 | 1,403 | 1,053 |
| total contig length (Gb) | 1.07 | 1.182 | 1.056 | 1.047 |
| maximum contig length (Mb) | 0.264 | 28.519 | 65.778 | 29.008 |
| contig N50 (Mb) | 0.026 | 5.534 | 17.655 | 4.378 |
| #scaffolds | 78,487 | 755 | 525 | 205 |
| longest scaffold length (Mb) | 5.998 | 207.238 | 197.608 | 151.897 |
| scaffold N50 (Mb) | 1.234 | 76.269 | 82.53 | 70.879 |
| total gap length (Mb) | 35.08 | 4.378 | 9.784 | 21.569 |
| anchored into chromosomes (%) | 25.9 | 95.6 | 98.6 | 97.2 |
| gap content (%) | 3.17 | 0.37 | 0.92 | 2.02 |
| BUSCO (%) | 91.5 | 94.2 | 95.1 | 95.1 |

## Analysis

### A much improved female duck genome

The final polished assembly (ZJU1.0) by Illumina reads exhibits a 62-fold improvement of scaffold continuity (N50 76.3Mb vs. 1.2Mb) compared to the Illumina genome, and is completely consistent with the FISH linkage map previously generated from 155 BAC clones (**Supplementary Fig. S2**)[33, 42]. The entire chrZ exhibits uniformly a 2-fold elevation of Illumina DNA sequencing read coverage in male relative to female, except for the chromosome tip of pseudoautosomal regions (PAR) (see below), confirming that we assembled the Z chromosome and that it does not have chimeric sequences with chrW or the autosomes. This new genome has 95.6% (1.13 Gb) of the assembled sequences assigned to 31 autosomes and the ZW sex chromosomes (**Supplementary Table S3**). The remaining 4.4% (62.1 Mb) of the genome not anchored or about 200Mb unassembled sequences based on the estimated genome size is likely due to their repetitive sequence composition or lack of linkage markers. In particular, the assembled macrochromosomes have become much more continuous (Figure 1b-c), and we have assembled majorities of microchromosomes that were all unmapped in the BGI1.0 genome (Figure 2a).

The ZJU1.0 genome assembly also has a higher level of completeness measured by its almost gapless sequence composition (0.37% vs. 3.17%), and substantial numbers of annotated telomeric and centromeric regions (Figure 2a, **Supplementary Table S4-5**), compared to the BGI1.0 assembly. We filled in a total of 116.2 Mb sequences of gaps within or between the BGI1.0 scaffolds, which were enriched for repetitive elements and GC-rich sequences (**Supplementary Fig. S3-4**). This can be explained by the inability of Illumina reads to span or resolve the repeat regions with high copy numbers or complex structures, and the sequencing bias against the GC-rich regions[43–45]. Indeed, we found specific transposable elements (TE) that are enriched in the filled gaps (**Supplementary Fig. S4**). These include the chicken repeat 1 (CR1) retroposon CR1-J2_Pass and the long terminal repeat (LTR) GGLTR8B that have undergone recent lineage-specific bursts in duck after its divergence with other Galloanserae species (Figure 2b, **Supplementary Table S6**). These apparent evolutionarily young repeats relative to other repeats of the same family in ducks show a lower level of sequence divergence from their consensus sequences (**Supplementary Fig. S5**), and tend to insert into other older TEs and form a nested repeat structure (**Supplementary Fig. S6**).

**Figure 2.**
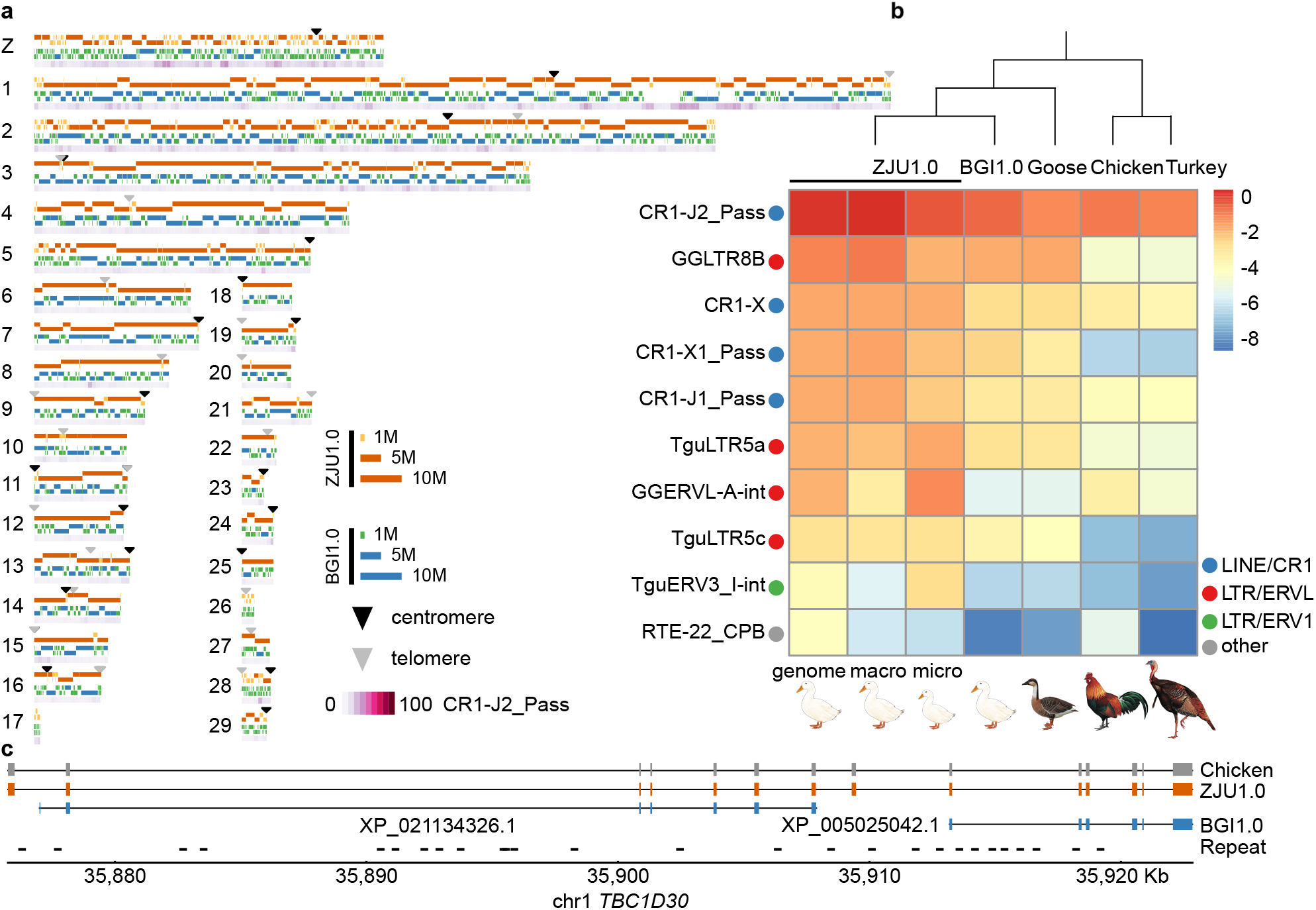
Comparing the new duck genome to other avian genomes. **a**. Schematic plot of each chromosome, showing the mapped contigs of ZJU1.0 (orange/yellow) and BGI1.0 (blue/green), putative centromeres (black triangles), and telomeres or interstitial telomeric sequences (grey triangles), and the most abundant repeat CR1-J2_Pass present in the gap regions of BGI1.0 (purple gradient). **b.** Comparisons of the top 10 most abundant repeats in the duck genome (ZJU1.0 whole genome, macrochromosomes, microchromosomes, and BGI1.0 assembly) to other Galloanseriformes bird genomes (goose, chicken, turkey). The more red, the higher proportion of assembled repeat content. **c.** An example gene annotation improvement showing two genes in the BGI1.0 genome are really one gene in the ZJU1.0 genome, and were fragmented into two because of low resolution of repeat sequences disrupting the previous genome assembly of exons.

Assembly of exon sequences embedded in such complex repetitive regions also led to the improvement of gene model annotations in our new assembly (e.g., Figure 2c). Overall, our new gene annotation combining a total of 17 duck tissue transcriptomes and chicken protein queries has predicted 15,463 protein-coding genes, including 71 newly annotated chrW genes. We have identified 8,238 missing exons in the BGI1.0 assembly in 2,099 genes, including 745 genes that were completely missing. We also corrected 683 partial genes, and merged them into 356 genes in the new assembly. The overall quality of our new duck genome is better than that of the previous Sanger-based zebra finch, and comparable to the latest version of chicken[41] and VGP zebra finch genomes[29] (Table 1).

### Different genomic landscapes of duck micro- and macrochromosomes

Our high-quality genome assembly and annotation of Pekin duck uncovered a different genomic landscape between the macro- and microchromosomes. Duck microchromosomes have a higher gene density than macrochromosomes per Mb sequence or per TAD domain (*P*< 2.2e-16, Wilcoxon test). The recombination rate estimated from the published population genetic data[46] is also on average 2.3-fold higher on microchromosomes than on macrochromosomes (16.3 vs. 7.2 per 50kb, *P*<2.2e-16, Wilcoxon test), which drives more frequent GC-biased gene conversion (gBGC) on the microchromosomes[47]. Both factors have resulted in a higher average GC content of the microchromosomes (Figure 3a-b; 44.5 % vs. 39.3 % per 50kb, *P*< 2.2e-16, Wilcoxon test). In addition, all chromosomes but chrZ (Figure 3a) show generally equal expression levels between sexes; genes on chrZ are expressed twice the level in males versus females. These chromosome-wide patterns are consistent with those reported in other birds regarding the differences between micro- and macrochromosomes, and a lack of global dosage compensation on avian sex chromosomes[1, 48, 49].

**Figure 3.**
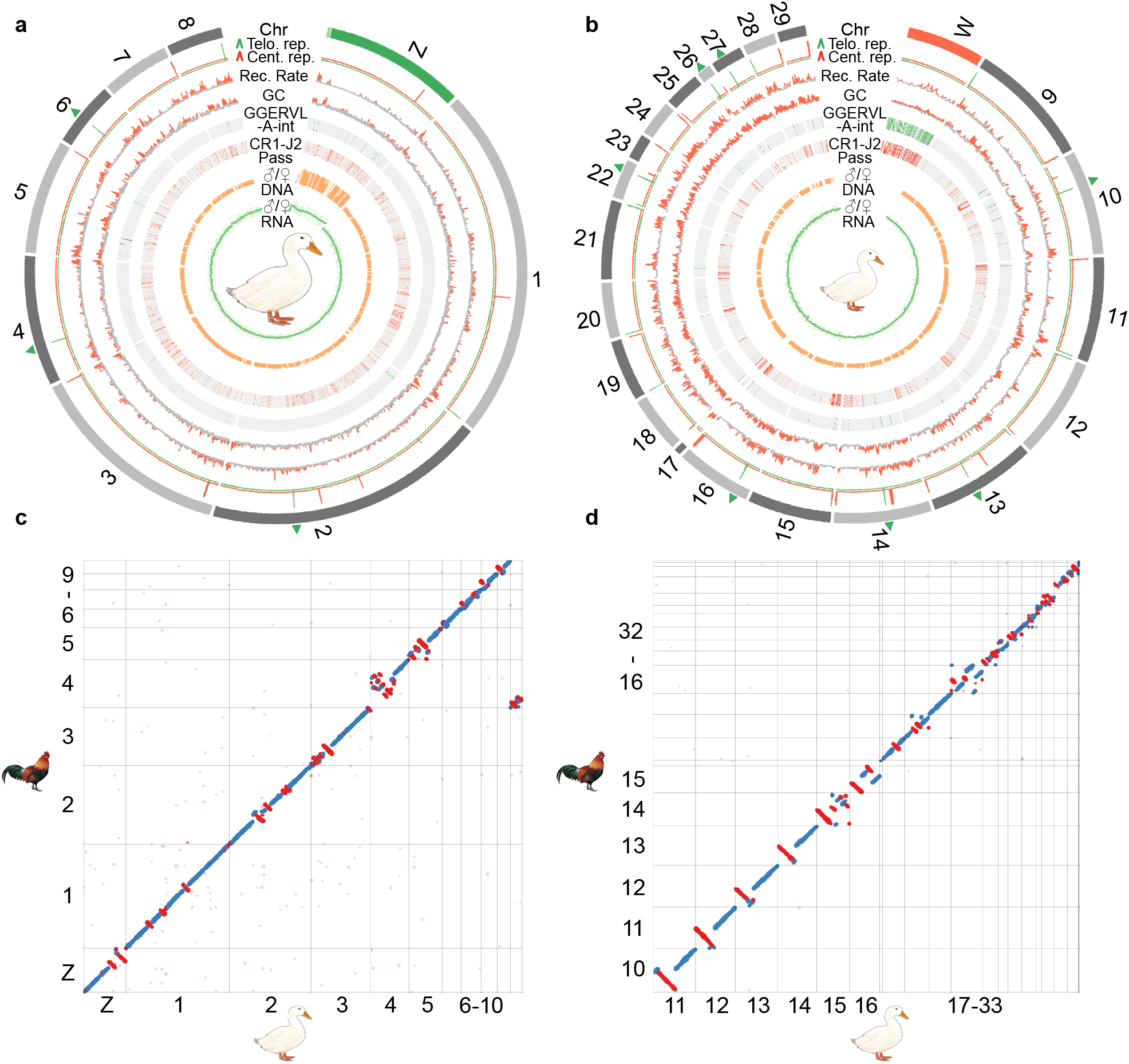
Evolution of the duck macro- and microchromosomes. From the outer to inner rings: the macro- (**a**) and microchromosomes (**b**), together with Z/W chromosomes (green/red color), and the pseudoautosomal regions (PARs) labelled with light green color at the tip of chrZ. Interstitial telomere sequences were labelled with green triangles on the chromosome. Putative centromeres (red lines) and telomeres (green lines) were inferred by the enrichment of centromeric and telomeric repeat copies, which show a sharp peak. We then show the recombination rate and GC content calculated in non-overlapping 50kb windows, as well as two repeat families (GGERVL-A-int and CR1-J2 Pass) that we identified to be enriched at centromeric regions and chrW. We also show the male vs. female (M/F) ratios of Illumina DNA sequencing coverage in non-overlapping 50kb windows, M/F expression ratios (each green dot as one gene) of the adult brain tissue and the smoothed line. **c-d.** Dot plots show the inversions between chicken and duck genome for both macro and micro chromosomes.

The completeness of our new duck genome is also demonstrated by its assembled centromeres (average length 443.3 kb) and telomeres (average length 73.7 kb), which were annotated by a cytogenetically verified *Anseriformes* centromeric repeat (ΛPL-*Hae*III)|50| and conserved telomeric motif sequences (**Supplementary Table S4-5**). We found 22 telomeric sites among the 31 chromosomes, of which 11 were interstitial telomeric repeat (ITR) sites inside the chromosomes (Figure 3a-b, green arrow heads). Consistent with the reported karyotypes of duck and other birds[50, 51], almost all microchromosomes are acrocentric indicated by their positions of centromeric region. Both macro- and microchromosomes centromeres are enriched for CR1-J2_Pass repeats (**Supplementary Fig. S7**), but microchromosome centromeres are specifically enriched for the LTR repeat GGERVL-A-int (Figure 3b, **Supplementary Fig. S8**). Such an interchromosomal difference of centromeric repeats has been reported in other birds and reptiles[52, 53], and is hypothesized to constitute the genomic basis for the spatial segregation of microchromosomes vs. machrochromosomes respectively in the interior vs. peripheral territories of the nucleus[54, 55]. Given their more aggregated spatial organization in the nuclear interior, microchromosomes exhibit an unusual pattern of more frequent inter-chromosomal interactions measured by the Hi-C data compared to macrochromosomes (**Supplementary Fig. S9**), consistent with the reported pattern of microchromosomes of chicken and snakes[56, 57].

To examine whether the different genomic landscape between micro-vs. macrochromosomes would underlie different frequencies or molecular mechanisms of intragenomic rearrangements during evolution, we used our newly produced chromosomal genome of emu (with a similar assembly pipeline to be reported in a companion paper[57]) as the outgroup, and identified 80 inversions on 26 chromosomes (>10kb, median size 1.5Mb, **Supplementary Table S7**) that occurred in the duck or *Anseriformes* lineage after it diverged from chicken in the past 72.5 MY[34] (Figure 3c-d). The average inversion rate (1.1 inversion events or 3.1Mb inverted regions per MY) of Pekin duck is lower than that of 1.5-2.0 events or 6.6-7.5Mb per MY between flycatcher and zebra finch[12], reflecting more frequent intragenomic rearrangements in the passerines[58, 59]. There are 46 inversions on the duck macrochromosomes, and 34 inversions on the microchromosomes, translating to 0.63 and 0.47 inversion events per MY, or 1.96 and 1.09 Mb inverted sequence per MY, respectively. A lower rate and shorter spanned length of inversions on the microchromosomes is probably related to their higher densities of genes and CNEs[60], because of the natural selection against inversions that disrupt these functional elements. Indeed, previous studies examining the breakpoint regions of genomic rearrangements of birds and mammals found that they tend to be devoid of CNEs[5, 61–63]. We also found that different families of TEs are significantly *(P*< 2.2e-16) enriched at the inversion breakpoints of macro-vs. microchromosomes relative to other genomic regions (**Supplementary Fig. S10**), suggesting they play an important role in mediating the inversions. However, we did not find a higher recombination rate at the breakpoint regions (**Supplementary Fig. S11**), unlike that reported previously in flycatcher and zebra finch[12, 15].

### Comparative analyses of topological chromatin domain architectures

Chromosomal inversions have attracted great interests of evolutionary biologists because they play an important role in local adaptation, speciation and sex chromosome formation[64]. We found that the duck or *Anseriformes* specific inversions (Figure 3c-d) are enriched for genes that function in immunity-related pathways (Figure 4a, e.g., ‘defense response to virus’, ‘G-protein coupled receptor pathway’; *P*<0.0001, Fisher’s Exact test), which may account for the known divergent susceptibility between chicken and duck against avian influenza virus. Indeed, RNF135 located on chr19, one of the ubiquitin ligases that regulate the RIG-I pathway responsible for the avian influenza virus response in ducks[65], is located in a duck-specific inversion.

**Figure 4.**
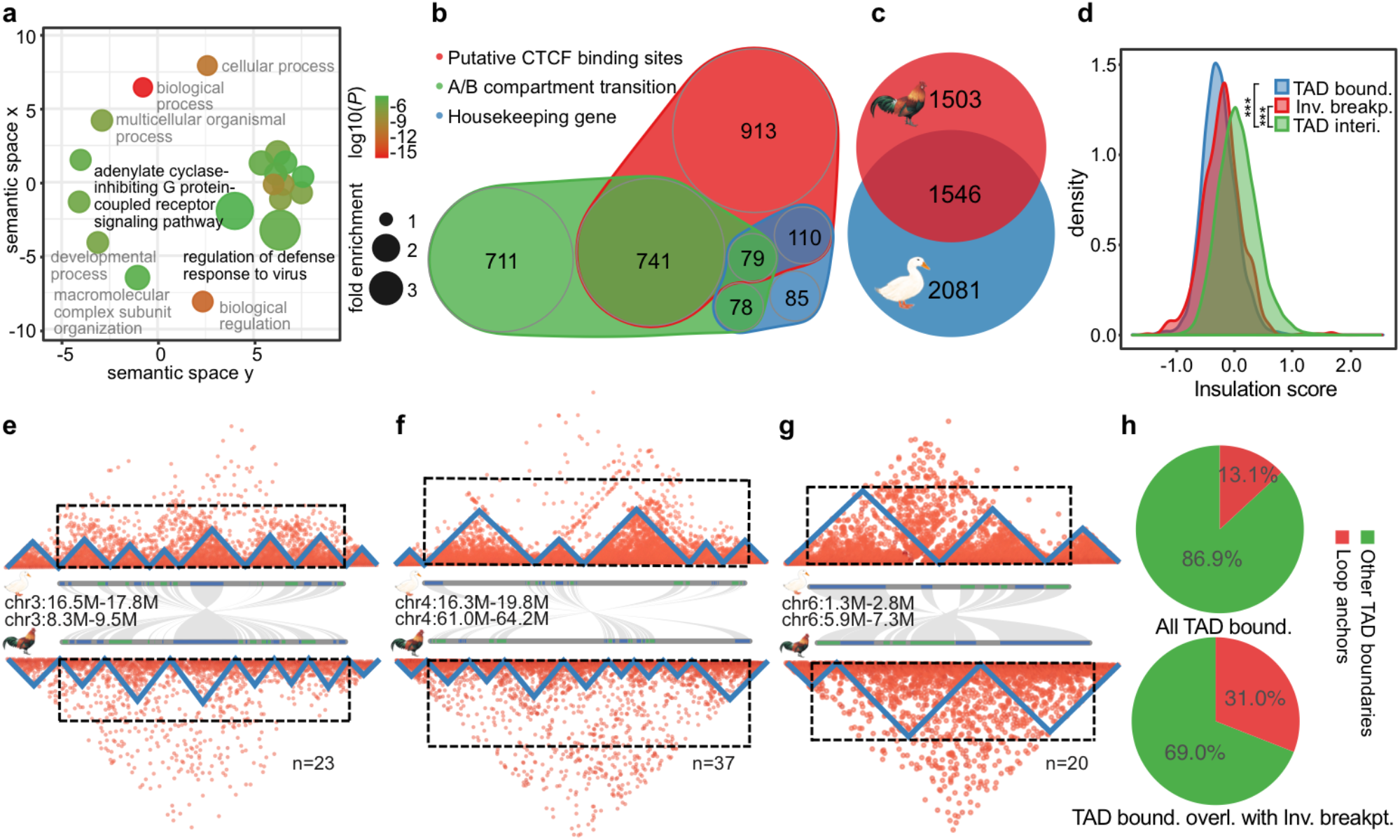
Genome inversions and topologically associated domains. **a**. Enriched GO terms of the genes included in the duck specific inversions. The x- and y-axes measure the GO term semantic similarities, which are used to remove the GO redundancies. **b**. Scaled Venn diagram shows the different compositions of TAD boundaries in duck. **c.** Scaled Venn diagram shows the TAD boundaries shared between chicken and duck. **d.** Inversion breakpoint regions tend to show a significantly lower insulation score than the TAD interior regions. **e-g**. We show the Hi-C heatmaps with each triangle structure indicating one TAD, along with the gene (blue or green bars) synteny plot between chicken and duck. Three examples are presented to show the impact of inversions between duck and chicken on TAD structure, with both inversion breakpoints (e), one inversion breakpoint (f), and no breakpoint (g), overlapped with the TAD boundaries. We also show the numbers of inversions that fit into each category. **h**. Pie charts showing that TAD boundaries that overlap with inversion breakpoints (bottom) have a higher percentage of loop anchors than others (top).

To systematically evaluate the functional impacts of the identified duck or *Anseriformes* specific inversions, we examined if there were any relationships with TAD units as well as their enclosed gene expression patterns compared to chicken. Similar to mammals[66], the boundaries of duck TADs are also characterized with a significant enrichment of putative binding sites of insulator protein CTCF (**Supplementary Fig. S12**), an enrichment of broadly expressed housekeeping genes (**Supplementary Fig. S13**), and coincide with the transitions between active (A) and inactive (B) chromatin compartments (**Supplementary Fig. S14**). The diverse types of TAD boundaries of duck are not mutually exclusive (Figure 4b), and suggest conserved mechanisms of TAD formation between birds and mammals[31]. The presence of putative CTCF binding sites, particularly with excessive pairs of binding sites in convergent orientation (‘loop anchors’) at the duck TAD boundaries (**Supplementary Fig. S15a-b**), suggested an active ‘loop extrusion’ mechanism involving both the extruding factors cohesin protein complex along chromatin and the counteracting CTCF protein[67]. In support of this, TAD boundaries that overlap with DNA loops have a significantly higher density of putative CTCF binding sites than any other TAD boundaries (**Supplementary Fig. S15c**). The overlap pattern between the TAD boundaries with the active/inactive compartment transition implies that self-organization of different chromatin types, probably driven by heterochromatin[68], underlies TAD formation. Finally, active transcription of genes[69] or TEs[70] have been recently discovered to account for TAD formation in mammals. We indeed found that various TEs located at the TAD boundaries have a significantly higher expression level (*P*<0.01, Wilcoxon test) than their copies elsewhere in the genome. However, these boundary TEs generally show a lower population frequency, and a higher level of segregating sequence polymorphism (*P*<0.05, Wilcoxon test) in their flanking sequences compared to the same families of TEs elsewhere (**Supplementary Fig. S16**), indicating that they are not under selection to fixation and may be recently inserted into the TAD boundaries. In addition, all the assembled centromere regions of metacentric chromosomes, and intriguingly 4 out of 11 ITRs (Figure 2a,b) coincide with the TAD boundaries (**Supplementary Figs. S7, 17**). This highlighted the uncharacterized role of ITRs in demarcating the functional domains in the chromosomes yet to be functionally tested in future.

We hypothesize that the TAD units or TAD boundaries are probably under strong selective constraint during evolution. This is suggested by some congenital diseases and cancer cases caused by disruptions of TADs through structural variations[71], and also sharing of TAD boundaries between distantly related species[66, 72]. A substantial proportion (42.6%) of duck TAD boundaries are shared with those of chicken (Figure 4c). This is probably an underestimate given that different tissues of Hi-C data were used here to identify TADs for the two bird species. A comparable level of conservation of human TAD boundaries (53.8%) has also been observed with mouse[66], and expectedly a lower level (26.8%) of conservation has been observed between human and chicken[56]. The other evidence of strong selective constraints acting on the integrity of TADs come from our findings here on the pattern of chromosomal inversion breakpoints of duck, whose TAD insulation scores are significantly (*P*< 2.2e-16, Wilcoxon test) lower (Figure 4d) than the TAD interior regions. That is, inversions more often precisely occurred at the TAD boundaries rather than within the TADs, i.e., disrupting the preexisting TADs. Only one third of the detected inversions have both their breakpoints located within the TADs, whereas the remaining two thirds have both or one of their breakpoints overlapping with the TAD boundaries (Figure 4e-g). Novel TAD boundaries that were created by the duck-specific inversions (e.g., Figure 4g) tend to have significantly higher insulation scores, i.e., weaker insulation strengths than those that are conserved between duck and chicken (**Supplementary Fig. S18**). This suggests that natural selection may more frequently target evolutionarily older and stronger TAD boundaries. We have to point out the alternative explanation for the overlap between the TAD boundaries and inversion breakpoints (Figure 4e) is that chromatin loop anchors bound by CTCF protein are more likely genomic fragile sites vulnerable for DNA double-strand breaks[73] that induce the inversions. Consistent with this explanation, we found that the TAD boundaries that overlap with inversion breakpoints (Figure 4h, **bottom**) have a significantly (*P*<0.001, Chi-square test) higher percentage of loop anchors than others (Figure 4h, **top**).

Since the novel TADs generated by chromosome inversions (e.g., Figure 4g) may create aberrant or new promoter-enhancer contacts, and consequently divergent gene expression during evolution, we further compared the levels of gene expression divergence in the conserved TADs vs. those novel TADs that encompass inversion breakpoints between chicken and duck. Interestingly, genes that are close to the novel TAD boundaries created by inversions only show slightly but not significantly higher levels of expression divergence than the genes located in the conserved TADs, except for certain tissues (**Supplementary Fig. S19**). This reflects that the TAD boundary changes have only affected a few genes’ expression patterns. It can be also explained by other regulatory divergences (e.g., in *cis*-elements) within the conserved TADs during the long-term divergence between chicken and duck, that have increased the target genes’ expression divergence to the same degree as that in the novel TADs.

### Sex chromosome evolution of Pekin duck

The Pekin duck provides a great model for understanding the process of avian sex chromosome evolution because the differentiation degree of its sex chromosomes is between those of ratites and chicken[27]. Previous comparative cytogenetic work found that the FISH probe of chicken chrZ cannot produce hybridization signals on chicken chrW because of their great sequence divergence, but instead can paint the entire chrW of duck and ostrich, suggesting that substantial sequence homology has been preserved between the Z/W chromosomes of the two species since the recombination was suppressed[27, 66]. The size of duck chrW is nevertheless smaller (estimated size 51Mb)[74, 75] compared to chrZ, probably because of extensive large deletions. Our new duck genome has assembled most of its chrZ derived from 53 scaffolds, except for 1.3 Mb unanchored sequences, into one continuous sequence 84.5Mb long (**Supplementary Fig. S20**). The size of duck chrZ is similar to that of published chicken chrZ (82.5 Mb[76]).

We determined 2.2Mb long PAR at the tip of chrZ (Figure 5a), based on its equal read coverage between sexes. This is consistent with previous cytogenetic work showing only one recombination nodule concentrated at the tip of the female duck sex chromosomes[77]. Consistently, the PAR shows a significantly (*P*< 2.2e-16, Wilcoxon test) higher rate of recombination than the rest Z-linked SDR that do not have recombination in females (Figure 5a). The distribution of GC content also exhibits a sharp shift at the PAR boundary because of the effect of gBGC (**Supplementary Fig. S21**). The evolution of chicken chrZ is marked by the acquisition of large tandem arrays of four gene families that are specifically expressed in testis[18]. In contrast, we did not find similar tandem arrays of testis genes on chrZ of duck, and all of the four Z-linked chicken testis gene families are located on the autosomes of duck (**Supplementary Fig. S22**).

**Figure 5.**
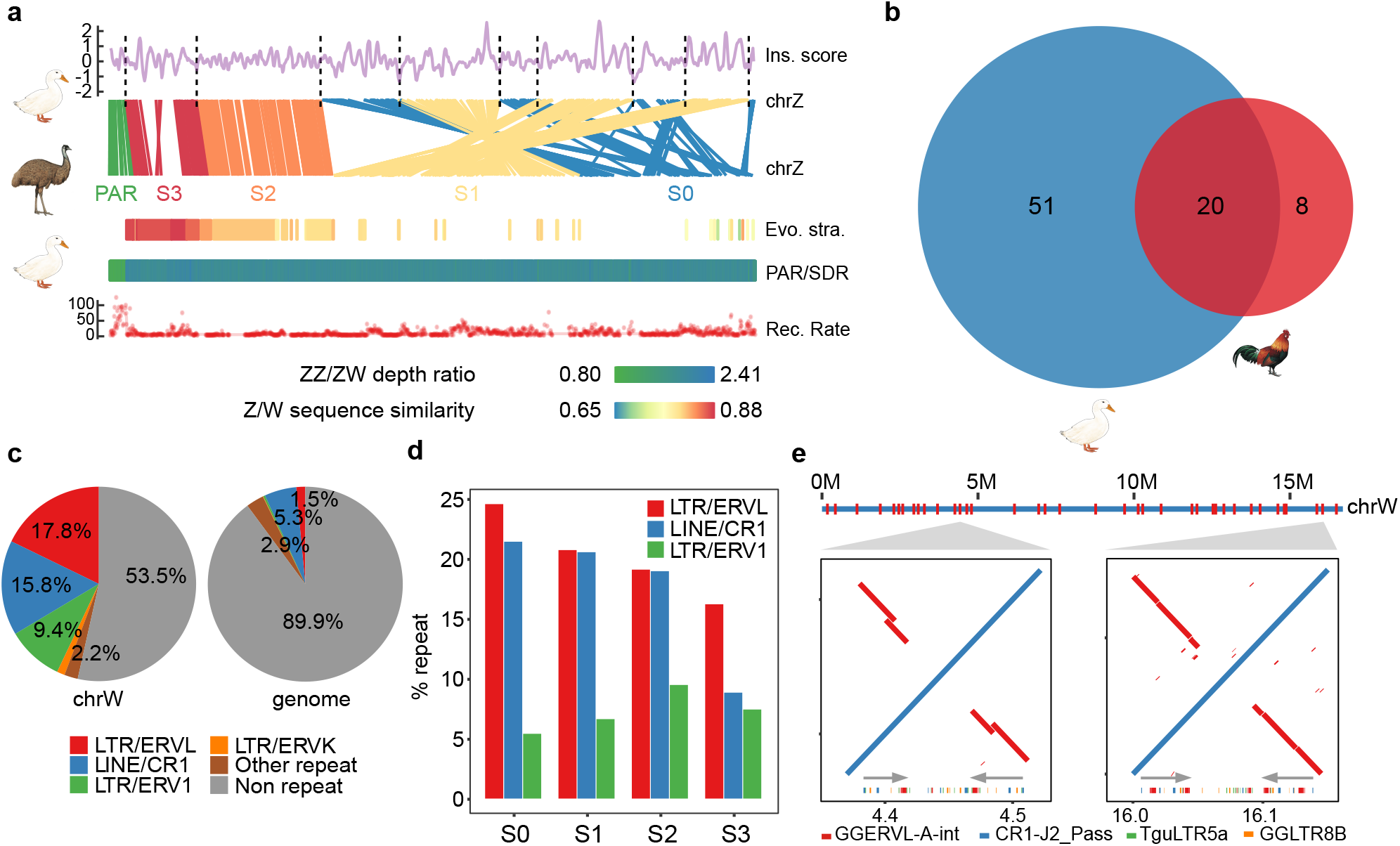
Sex chromosome evolution in Pekin duck. **a**, Evolutionary strata analyses of the duck sex chromosomes. From top to bottom: the breakpoints of genomic rearrangements between emu and duck chrZ tend to have a lower insulation score; gene synteny between the emu and duck Z chromosomes; alignment of the duck chrW scaffolds against the emu chrZ reveals a pattern of evolutionary strata, with each scaffold showing the color-scaled sequence divergence levels between the duck chrW vs. the emu chrZ; PAR (light green)/SDR (dark green) composition inferred by the ratio of male vs. female Illumina DNA sequencing depth with the color scaled to the ratio value; a higher recombination rate in the duck PAR than in SDR. **b**. Scaled Venn diagram showed the chrW genes shared between duck and chicken. **c.** Comparing the repeat content of the duck chrW to the whole genome. **d**. Different enrichment trends of chrW repeats at different evolutionary strata. **e**, Palindrome structure of duck chrW. Palindromes are labelled across the entire chrW (red), ordered according to the duck chrZ. Shown are alignment plots of two zoomed-in examples of palindromes (red inversions and grey arrows) for their repeat content (colors below grey arrows).

The assembled duck chrW assembly contains 36 scaffolds with a total length of 16.7Mb (about one third of the estimated size), all of which are almost exclusively mapped by female reads (**Supplementary Fig. S20**). It marks an 8.8-fold increase in size compared to our previous assembly using Illumina reads[20, 78], and is much longer than the most recent assembly of chicken chrW (6.7 Mb)[22]. We have annotated a total of 71 duck W-linked SDR genes, and all of them are single copy genes, compared to 27 single-copy genes and one multicopy gene on the chicken chrW, with 20 genes overlapped between the two (Figure 5b). The only multicopy chicken W-linked gene *HINTW* with about 40 copies[22] is present as a single-copy gene on the duck chrW. These results indicate that duck and chicken have independently evolved their sex-linked gene repertoire since their species divergence. The duck chrW retained more genes than chicken, and represents an intermediate stage of avian sex chromosome evolution between those of ratites and chicken.

Due to the intrachromosomal rearrangements of chrZ, most birds (including duck) except for ratites have retained few ancestral gene syntenies of their proto-sex chromosomes before the suppression of homologous recombination[20, 78], and exhibit dramatic reshuffling of their old evolutionary strata. In order to accurately reconstruct the history of duck sex chromosome evolution, we used a newly produced chrZ assembly of emu in our group to approximate the avian proto-sex chromosomes. Almost all (15.2Mb, 91%) of the duck chrW sequences can be aligned to the chrZ of emu, and form a clear pattern of four evolutionary strata. This is manifested as a gradient of Z/W pairwise sequence divergence, i.e., a gradient of the age of strata along the chrZ, which is named from the old to the young, as stratum 0, S0 to S3, (Figure 5a). Within each stratum, chrW scaffolds of similar levels of sequence divergence are clustered and separated from the neighbouring strata with different divergence levels (**Supplementary Fig. S23**). The genes enclosed in each stratum are consistent with our previous annotation of the duck evolutionary strata based on the BGI1.0 genome, and show a consistent gradient of synonymous substitution rates (**Supplementary Fig. S24**) between the Z-and W-linked alleles according to the age of the strata where they reside. We did not find any chrW scaffolds that span the boundaries of neighbouring strata, probably because of some complex repeat sequences (e.g., CR1-J2_Pass) that accumulate at the boundary. Interestingly, the inferred boundaries between evolutionary strata on chrZ, i.e., the breakpoints between the inverted regions within or between the strata (8 out of 9 boundaries shown in Figure 5a) tend to have a low TAD insulation score, i.e., to overlap with TAD boundaries or loop anchors (**Supplementary Fig. S25**). This again strongly supports the idea that loop anchors or TAD boundaries are likely the genomic fragile regions that induced inversions.

Because of the lack of recombination, majorities (30 or 42.9%) of W-linked genes probably have become pseudogenes or long non-coding RNA genes due to frameshift mutations or premature stop codons (**Supplementary Fig. S26**). The other pronounced signature of functional degeneration of chrW is accumulation of TEs. The duck chrW shows a much higher genomic proportion (46.5% vs. 10.1%) and a different composition of TEs compared to the genome average (Figure 5c). The W-linked repeats are concentrated in those families that have specifically expanded their copy numbers in the duck after it diverged from other *Anseriformes* (**Supplementary Fig. S27, Supplementary Table S8**). Among them, different TE families exhibit opposing trends of colonizing the different evolutionary strata of different ages (Figure 5d, **Supplementary Fig. S28**). TE families that have been propagating since the ancestor of Neoaves (e.g., CR1-J2_Pass, **Supplementary Fig. S6**)[79] are more enriched in the older strata, while TE families that were specifically propagated in the duck (e.g., TguERV3_I-int, Figure 2b) are more enriched in the younger strata. This suggests that older evolutionary strata might be saturated for old TEs relative to TEs with recent activities. Particularly, duck or *Anseriformes* enriched repeats are nested with each other and form 38 palindromes dispersed across the entire chrW (Figure 5e). Their lengths range from 15.2 kb to 345.5 kb (**Supplementary Table S9**), together comprising 3.74Mb or 22% of the assembled duck chrW sequence.

## Discussion

Birds and mammals diverged over 300 MY ago and are known to have a very different chromosomal composition[1]. Our comparative analyses of the nearly complete genome of the Pekin duck revealed that TADs are conserved functional and evolutionary chromosome units in both birds and mammals. The 40% to 50% of the TADs shared between chicken and duck is comparable to the proportions shared between human and mouse[66]. This is also consistent with the highly conserved pattern of replication domains between human and mouse[80], which have a nearly one-to-one correspondence with TADs[81]. The interspecific overlap of TADs implies strong selection on TAD integrity during evolution. In this work, we identified many chromosomal inversions between chicken and duck that were previously uncharacterized because of the fragmented duck Illumina-based genome. Consistent with selection against the genome rearrangements disrupting the TADs, there are disproportionately more chromosome inversions that occurred at the TAD boundaries than within the TADs. This extensive overlap between TAD boundaries and inversion breakpoints likely reflects the susceptibility of TAD boundaries to DNA double-strand breaks. TADs can form either by self-organization of genomic regions of the same epigenetic state, or by active loop extrusion involving the cohesin and insulator protein CTCF[67]. This is indicated by the transition between active and inactive chromatin compartments or the enrichment of CTCF binding sites at the TAD boundaries of duck (this study), chicken[56], and mammals[66]. It has been recently shown that type II topoisomerase B (TOP2B), which releases the DNA torsional stress by transiently breaking and rejoining DNA double-strands, physically interacts with cohesin and CTCF and colocalizes with the TAD boundaries with convergent CTCF binding site pairs (loop anchors)[73]. This probably frequently exposes the TAD boundaries to double-strand breaks, and induces chromosomal inversions involving the entire TAD. This mechanism may also account for the common genomic fragile sites found in both birds and mammals that have been reused during evolution to mediate genomic rearrangements[7, 11, 13, 82]. Overall, despite divergent chromosomal composition, our results suggested conserved mechanisms of chromosome folding and rearrangements between birds and mammals.

The two clades of vertebrates also evolved convergent sex chromosome architectures. Our finding that the duck chrW has suppressed recombination with chrZ in a stepwise manner is similar to the pattern of evolutionary strata between the human X and Y chromosomes[19]. As the result of recombination suppression, the duck chrW has accumulated massive TEs, some of which formed dispersed palindromes along the chromosome. Unlike other sex-specific palindromes reported in primates, birds and willow[25, 26, 83–85], the duck palindromes do not seem to contain functional genes that have robust gene expression. This suggests that the gene copies contained in the palindromes may have nevertheless become pseudogenes, despite the repair mechanism mediated by gene conversions between gene copies within the palindromes. Or the involved genes have already become a pseudogene before being amplified by the palindromes. An interesting contrast is that we did not find palindromes on our recently assembled emu chrW with a similar dataset and pipeline, which evolves much slower than chrWs of chicken and duck. Palindromes were also not reported in the recently evolved *Drosophila miranda* chrY[86]. These results suggest that sex-linked palindromes are a feature of strongly differentiated sex chromosomes which have accumulated abundant TEs. The palindromes may retard the functional degeneration of Y- or W-linked genes, but can also promote large sequence deletions by intrachromosomal recombination. The latter probably contributed to the much smaller size of chrW relative to the chrZ of duck, despite many more genes than the chrW of chicken have been preserved.

## Methods

### Genome assembly

High molecular weight DNA (HMW DNA) was extracted from the liver of a female Pekin duck *(Anas platyrhynchos*, Z2 strain) with Gentra Puregene Tissue Kit (Qiagen #158667). Libraries for SMRT sequencing were constructed as described previously[87]. In total, 115 SMRT cells were sequenced with PacBio RS II and Sequel platform (Pacific Biosciences), and 186 Gb (143-X genome coverage) subreads with an N50 read length of 14,262 bp were produced. The same DNA was used to generate a linked-reads library following the protocol on the 10X Genomics Chromium platform (Genome Library Kit & Gel Bead Kit v2 PN-120258, Genome HT Library Kit & Gel Bead Kit v2 PN-120261, Genome Chip Kit v2 PN-120257, i7 Multiplex Kit PN-120262). This 10X library was subjected to MGISEQ-2000 platform for sequencing and 185 Gb PE150 (142-X genome coverage) reads were collected. HMW DNA of a male Pekin duck was used to produce the BioNano library with the Enzyme Nt.BspQ1. After the enzyme digestion, segments of the DNA molecules were labeled and counterstained following the IrysPrep Reagent Kit protocol (Bionano Genomics) as described previously[88]. Libraries were then loaded into IrysChips and run on the Irys imaging instrument, and a total of 73 Gb (56-X genome coverage) optical map data were generated. We used the HMW DNA from the breast muscle of a male Pekin duck to prepare the Hi-C library using the restriction enzyme Mbol with the protocol described previously[30] and produced a total of 106Gb (82-X genome coverage) pair-end reads of 50bp long on the Illumina HiSeq X Ten platform. We used the published genome resequencing data of 14 female and 11 male duck individuals from[46]. We collected the total RNAs of adult tissues (brain, kidney, gonads) of both sexes using TRIzol^®^ Reagent (Invitrogen #15596-018) following the manufacturers’ instructions. Then paired-end libraries were constructed using NEBNext^®^ UltraTM RNA Library Prep Kit for Illumina^®^ (NEB, USA) and 3Gb paired-end reads of 150bp were produced for each library.

We generated the genome assembly with the modified Vertebrate Genomes Project (VGP) (v1.0) pipeline[29]. In brief, we produced the contig sequences derived from the PacBio subreads using FALCON[89] (git 12072017) followed by two rounds of assembly polishing by Arrow[90], and then by Purge Haplotigs[91] (bitbucket 7.10.2018) to remove false haplotype and homotypic duplications. The contigs were then scaffolded first with 10x linked reads using Scaff10X (https://github.com/wtsihpag/Scaff10X), then with BioNano optical maps using runBNG[92] (v1.0.3), and finally with Hi-C reads using SALSA[93] (v2.0). We performed gap filling on the scaffolds with the Arrow-corrected PacBio subreads by PBJelly[94], and two rounds of assembly polishing with Illumina reads by Pilon[95] (v1.22). All the scripts used from the VGP assembly pipeline[29] are available at https://github.com/VGP/vgp-assembly. We evaluated the genome completeness using BUSCO[96] (v3.0.2). In brief, 4,915 benchmarking universal single-copy ortholog (BUSCO) proteins of birds from OrthoDB v9 were used in the evaluation.

### Genome annotation

We combined evidence of protein homology, transcriptome and *de novo* prediction to annotate the protein-coding genes. First, we aligned the protein sequences of human, chicken, duck and zebra finch collected from Ensembl[97] (release 90) to the reference genome using TBLASTN[98] (v2.2.26) with parameters: -F F -p tblastn -e 1e-5. The resulting candidate genes were then refined by GeneWise[99] (v2.4.1). For each candidate gene, only the one with the best score was kept as the representative model. We filtered the candidate genes, if they contain premature stop codons or frameshift mutations reported by GeneWise[99]; or if single-exon genes with a length shorter than 100bp, or multi-exon genes with a length shorter than 150bp; or if the repeat content of the CDS sequence is larger than 20%. Second, to obtain the *de novo* gene models, we used the protein queries to train Augustus[100] (v3.3) with default parameters. We also used all available RNA-seq reads to construct transcripts using Trinity[101] (v2.4.0). Finally, all the gene models from the above three resources were merged into a non-redundant gene set with EVidenceModeler[102] (v1.1.1). We used RepeatMasker[103] (v4.0.8) with parameters: -s -pa 4 -xsmall, and the RepBase[104] (v21.01) queries to annotate the repetitive elements.

To annotate the putative centromeres, we searched the genome with the reported 190bp duck centromeric repeats[50] using TRFinder[105] (v4.09) with the parameters: 2 5 7 80 10 50 2000. A genome-wide distribution of the 190bp sequences was generated by binning the genome with a 50kb non-overlapping window to find the local enrichment of copy numbers, which was defined as the putative centromeres. For telomeres, we used the known vertebrate consensus sequence[106] ‘TTAGGG/CCCTAA’ to search for the clusters of consensus sequence on both strands from the above tandem repeat annotation. Consensus sequence enriched genomic blocks in a 50kb window were then defined as the putative telomere regions.

### Building the chromosomal sequences and identifying the sex-linked sequences

To anchor Pekin duck scaffolds onto chromosomes, we first collected the ordered 1689 RHmap linked contigs[32] and 155 BAC clone sequences[33] from the previous studies. We aligned these sequences, as well as the Illumina duck genome[36] (BGI1.0) to the new duck scaffolds we generated by nucmer[107] (v3.23) packages (http://mummer.sourceforge.net) and only kept the best hits for each sequence. Scaffolds were orientated and ordered first based on the RHmap contigs that span more than one scaffold, then by BAC sequences whose order was determined previously by FISH, and finally by the syntenic relationship with the BGI1.0 genome. We also corrected scaffolding errors using the raw PacBio reads, if the order of our scaffolds had conflicts with that of RHmap or BAC sequence order (**Supplementary Fig. S2**).

To identify the sex-linked sequences, Illumina reads from both sexes were aligned to the scaffold sequences using BWA ALN[108] with default parameters. Read depth of each sex was then calculated using SAMtools[109] in 5kb non-overlapping windows, and normalized against the median value of depths per single base pair throughout the entire genome, respectively, to enable the comparison between sexes. To identify the Z-linked sequences, the depth ratio of male-vs- female (M/F) was calculated for the genomic regions mapped by reads for each sequences, with a minimum 80% coverage in both sexes, and sequences with a depth ratio ranging from 1.5 to 2.5 were assigned as Z-linked. To identify the W-linked sequences, we calculated M/F depth ratio as well as M/F coverage ratio and assigned scaffolds to W-linked when either ratio was within the range from 0.0 to 0.25 as W-linked sequences (**Supplementary Fig. S21**). Since we do not have linkage markers on the W chromosome, we ordered the W scaffolds based on their unique aligned position with the Z chromosome using RaGOO[110] (v1.1) with default parameters (https://github.com/malonge/RaGOO). This does not reflect the actual order of W-linked sequences which probably have rearrangements with the homologous Z chromosome, but allows us to examine the pattern of evolutionary strata.

To identify the inversions in the duck genome, genomic syntenic blocks between chicken and duck, and emu and duck were constructed using nucmer (v3.1) with the parameters: -b 500 -l 20. Then inversions between chicken and duck were manually checked by plotting the dot plot between the two species. The duck specific inversions were identified by excluding chicken-specific inversion, using emu as the outgroup.

### Hi-C analyses

Hi-C read mapping, filtering, correction, binning and normalization were performed by HiC-Pro[111] (v2.10.0) with the default parameters. In brief, Hi-C reads of chicken[112] (sourced from FR-AgENCODE project) and duck were mapped to the respective reference genome and only uniquely mapped reads were kept. Then each uniquely mapped reads were assigned to a restriction fragment and invalid ligation products were discarded. Data was then merged and binned to generate the genome-wide interaction maps at 10kb and 50kb resolution. TADs were identified by HiCExplorer[113] (v3.0) with the application hicFindTADs. First, HiC-Pro interaction maps were transformed to h5 format matrix by hicConvertFormat with parameters: --inputFormat hicpro --outputFormat h5. Then the h5 matrix was imported to hicFindTADs with parameters:--outPrefix TAD --numberOfProcessors 32 --correctForMultipleTesting fdr. hicFindTADs identifies the TAD boundaries through an approach that computes a TAD insulation score. Genomic bins with low insulation scores relative to neighboring regions were defined as local minima and called as the TAD boundaries. Human CTCF[114] motif was used as a query for FIMO in MEME[115] (v4.12.0) to identify the putative CTCF binding sites. CTCF density in every 10kb non-overlapping sliding window along the genome was calculated to check its enrichment at the TAD boundaries. We identified the A/B compartments using the pca.hic function from HiTC[116] (High Throughput Chromosome Conformation Capture analysis) R package with default parameters, and the 10kb matrix generated by HiC-Pro as the input. We identified the chromatin loops by Mustache[117] with the parameters: -p 32 -r 10kb -pt 0.05, after converting the h5 format matrix to mcool matrix format by hicConvertFormat with parameters: --inputFormat h5 --outputFormat mcool.

### Evolutionary strata

To demarcate the evolutionary strata, all the repeat masked duck W-linked scaffolds were aligned to emu Z chromosome using LASTZ[118] (v0.9) with parameters: --step=19 --hspthresh=2200 -- inner=2000 --ydrop=3400 --gappedthresh=10000 --format=axt, and a score matrix set for the distant species comparison. Alignments were converted into ‘net’ and ‘maf’ results using UCSC Genome Browser’s utilities (http://genomewiki.ucsc.edu/index.php/). Based on ‘net’ and ‘maf’ results, the identity of the aligned sequence was calculated for each alignment block with a 10kb nonoverlapped window and then we oriented the aligned W-linked sequences along the Z chromosomes. Then we color-coded the pairwise sequence divergence level between the Z/W sequences to demarcate the evolutionary strata.

### Gene expression analyses

RNA-seq reads were mapped to the duck genome by HISTA2[119] with default parameters. Only uniquely mapped RNA-seq reads were kept and used to calculate the RPKM expression level. DESeq2[120] was applied to normalize the RPKM values across different samples and finally generated an expression matrix. For each gene, we used the median expression value in each tissue to calculate the tissue specificity index TAU[121, 122]. Expression levels of TE elements were calculated using SQUIRE[123] (v0.9.9.92) (https://github.com/wyang17/SQuIRE) with default parameters.

## Data availability

The assembly and annotation of Pekin duck has been deposited in GenBank under the Bioproject accession code PRJNA636121 (accession number JACGAL000000000) and the emu under PRJNA638233 (accession number JABVCD000000000).

## Code availability

Scripts used in this study are shared on GitHub at https://github.com/ZhouQiLab/DuckGenome

## Acknowledgment

Q.Z. is supported by the National Natural Science Foundation of China (31722050, 31671319), the Natural Science Foundation of Zhejiang Province (LD19C190001) and the European Research Council Starting Grant (grant agreement 677696). We thank BGI-Shenzhen for providing the 10x linked reads data of duck.

## Conflict of interest statement

None declared.

## Authors’ contributions

Q. Z. conceived the project and acquired the funding; J. L., X. D., S. F., C. G., J. R., K. W., acquired the samples and produced the data; J. L., J. Z., J. L., Y. Z., C. C., L. X., Q. Z. performed the analyses.; J. L., Y. J., Z. Z., G. Z., E. J. and Q. Z. wrote the paper.

## Notes

### Competing Interest Statement

The authors have declared no competing interest.

