## Supplementary Figures for "A new duck genome reveals conserved and convergently evolved chromosome architectures of birds and mammals"

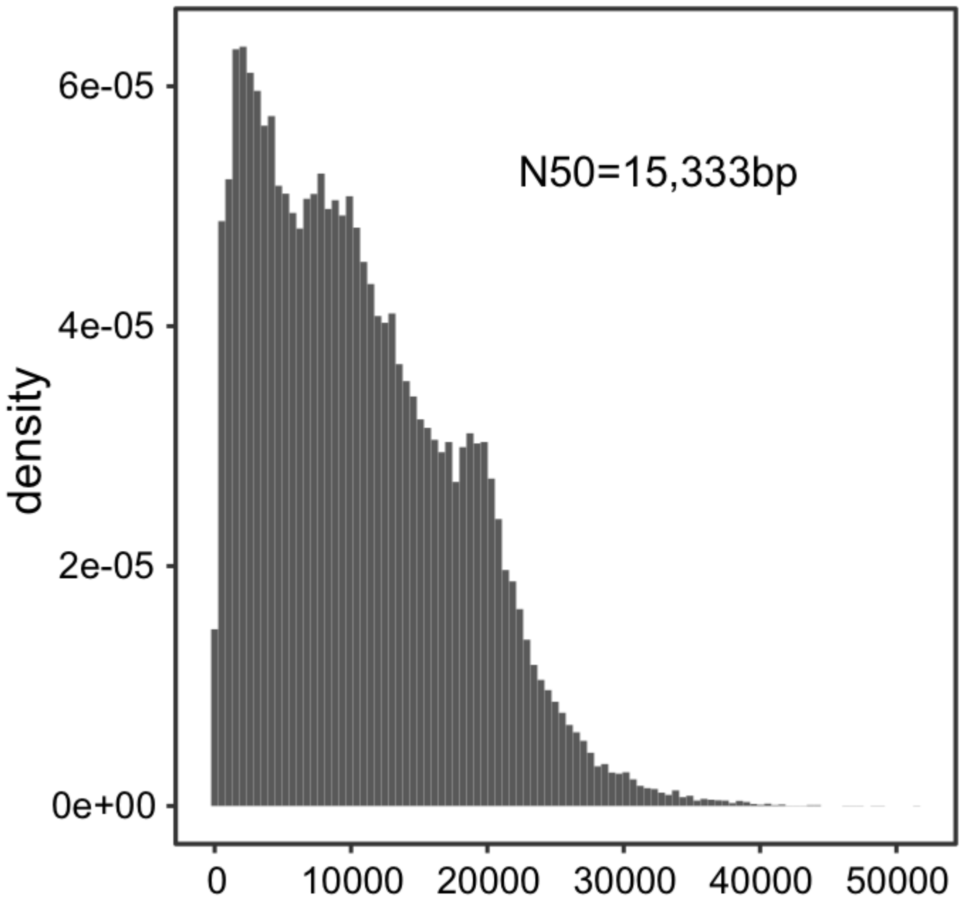


**Supplementary Figure S1 | Length distribution of one representative Pacbio RSII SMRT cell from all 115 SMRT cells.** The subread N50 length is 15,333 bp.

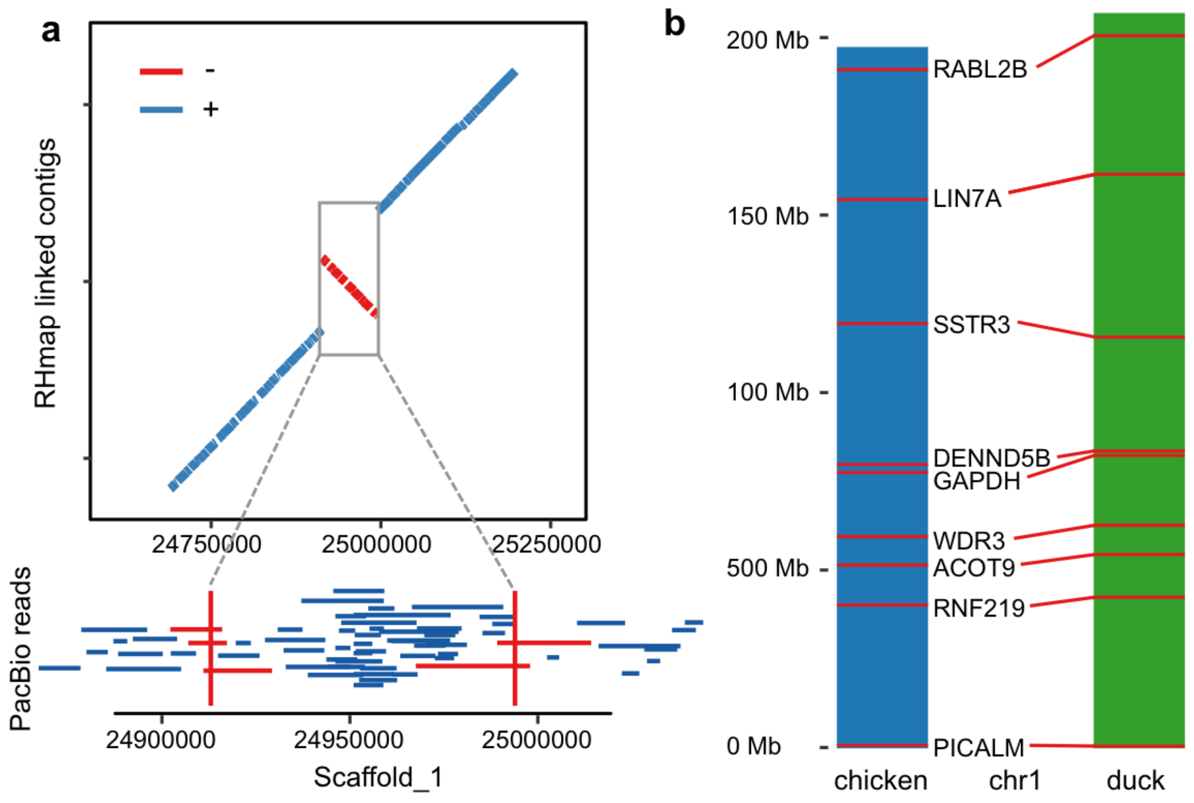


**Supplementary Figure S2 | a. A representative case of assembly error correction.** The conflict of orientation is seen as an inversion between the RH map (y axis) and the Hi-C scaffold (Scaffold_1; x-axis). To infer the correct orientation, we mapped the PacBio raw reads to the inversion breakpoints (red). In this case, the PacBio reads supported the RH map, and the assembly error was corrected. **b. A representative case to show the duck assembly is consistent with the FISH linkage map.** Here we used the chr1 as an example, the gene order is consistent between chicken and duck assembly.

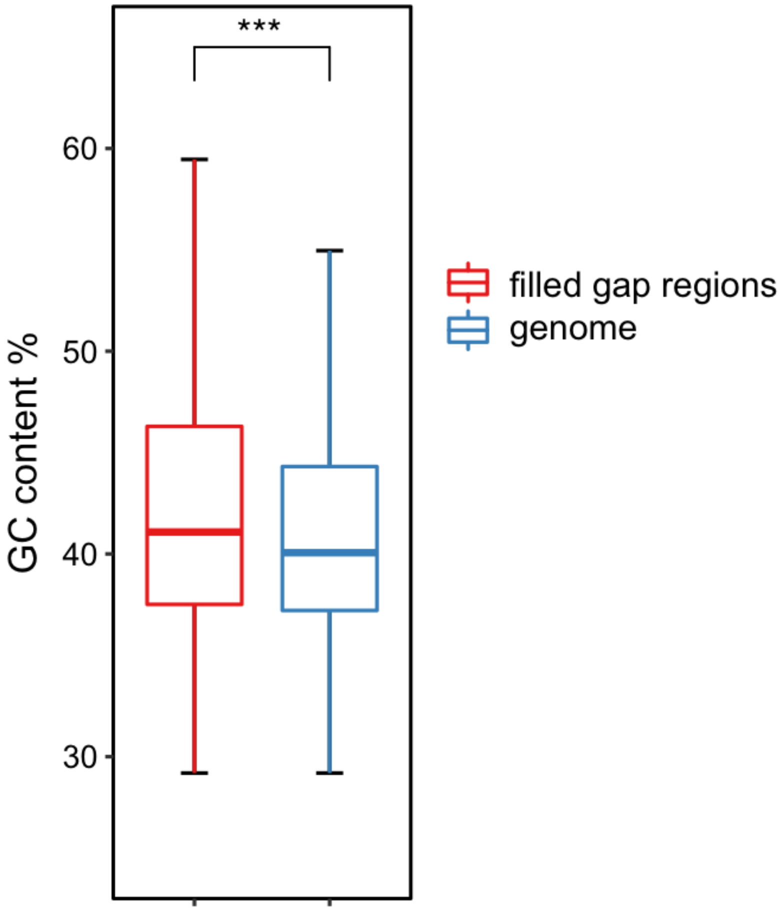


**Supplementary Figure S3 | High GC content at the gap regions of BGI1.0.** The ZJU1.0 genomic regions corresponding to the gap regions of BGI1.0 have a significantly higher GC content than the rest of the genome. “***” indicates P < 2.2e-16, measured by one-sided Wilcoxon signed rank test.

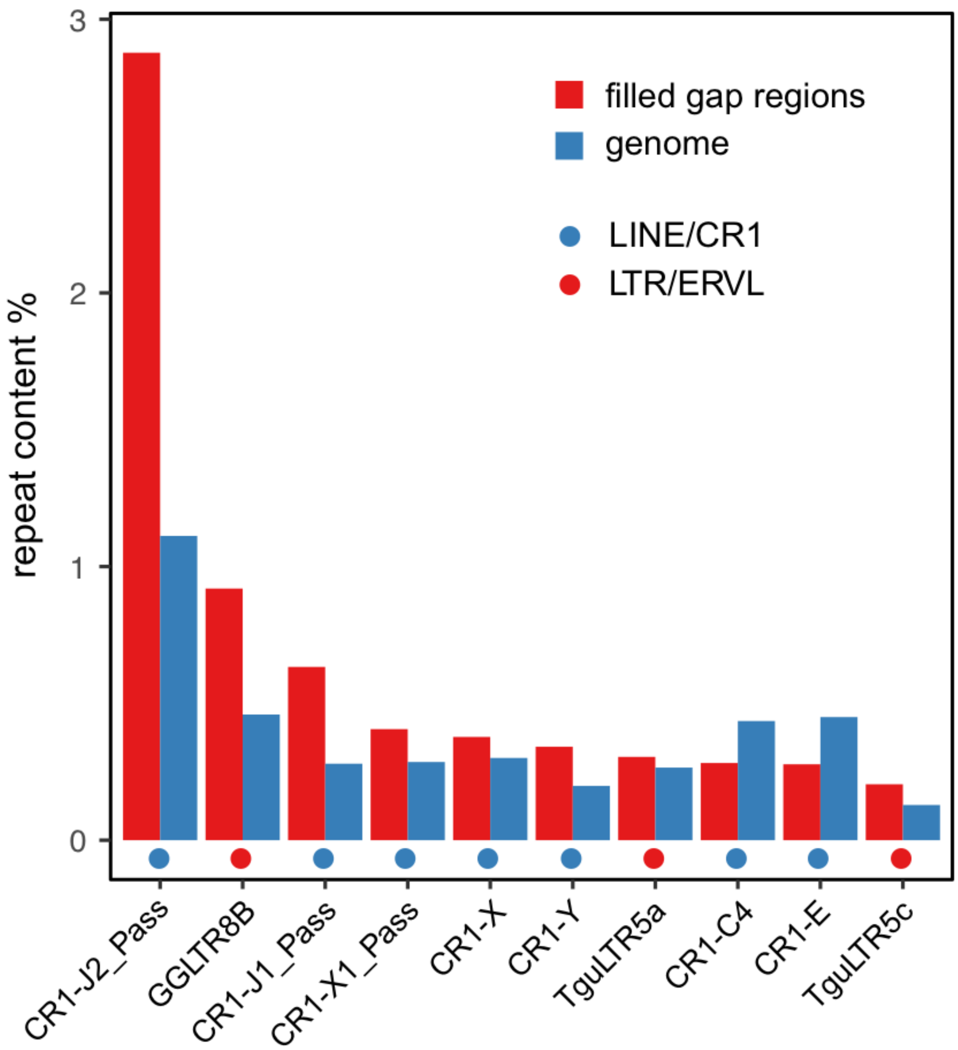


**Supplementary Figure S4 | Transposable elements (TE) are enriched in the filled gap regions.** Compared to the whole genome, the filled gap regions are mainly enriched for two kinds of repeats: LINE/CR1 and LTR/ERVL. The most abundant repeat in filled gap regions is CR1-J2_Pass.

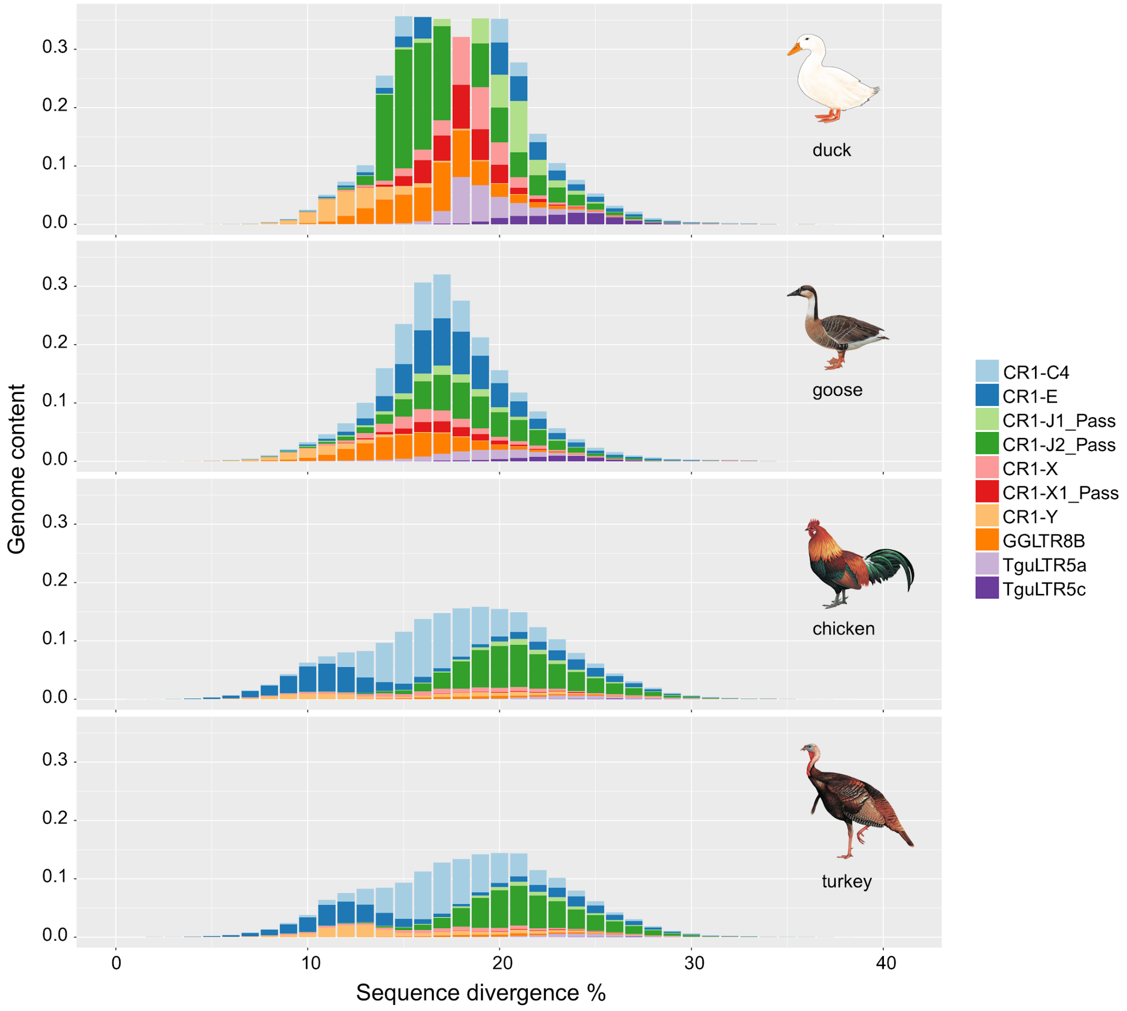


**Supplementary Figure S5 | Comparison of repeat composition between duck and other Galloanseriformes birds.** The repeats that are specifically enriched in the filled gap regions show a lower level of sequence divergence from their consensus sequences, i.e., tend to be young repeats.

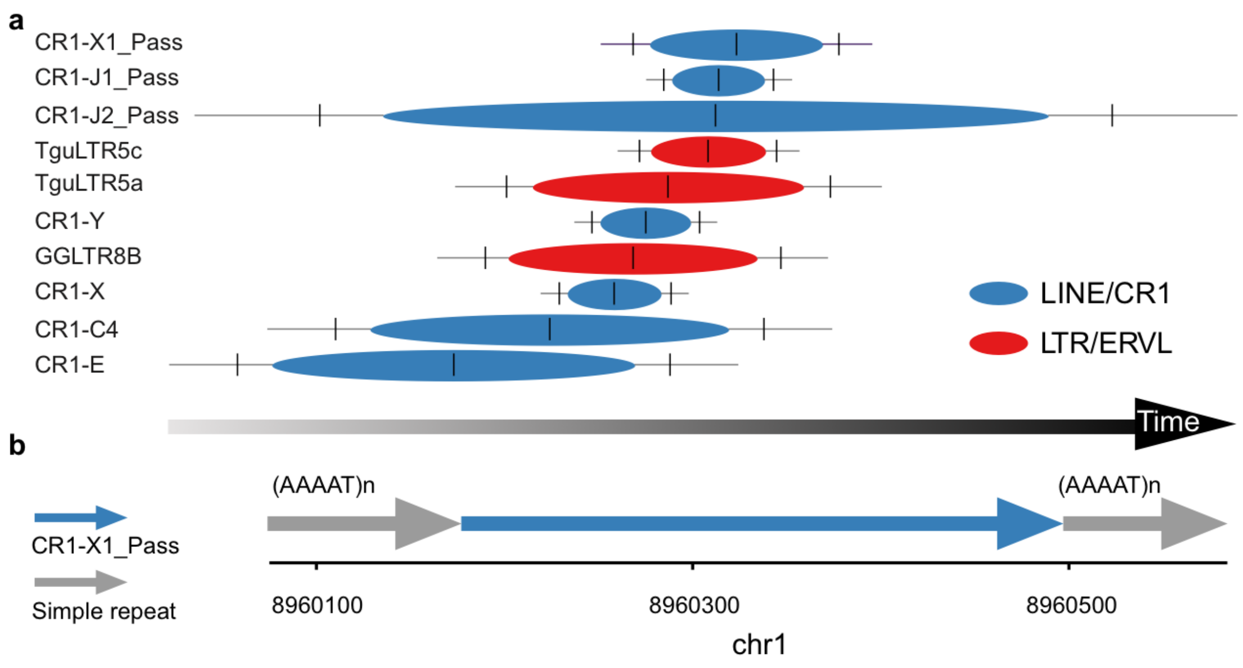


**Supplementary Figure S6 | Transposition in transposition (TinT) analyses of repeats enriched in the filled gap regions. a.** A graphic presentation to indicate the chronological activities of the top 10 enriched TEs in the BGI1.0 gap regions sorted by the peak of each activity. The center of each oval represents the peak of the activity period; the probable activity period ranges that cover 75%, 95% and 99% are indicated by the ends of each oval, the vertical lines and the ends of each line, respectively. Two classes of TE are colored in red (LINE/CR1) and blue (LTR/ERVL). From left to right indicates the inferred age of each repeat family from a more ancestral time point to a more recent time point. **b.** A case to show CR1-X1_Pass was inserted into one simple repeat and formed a nested repeat structure.

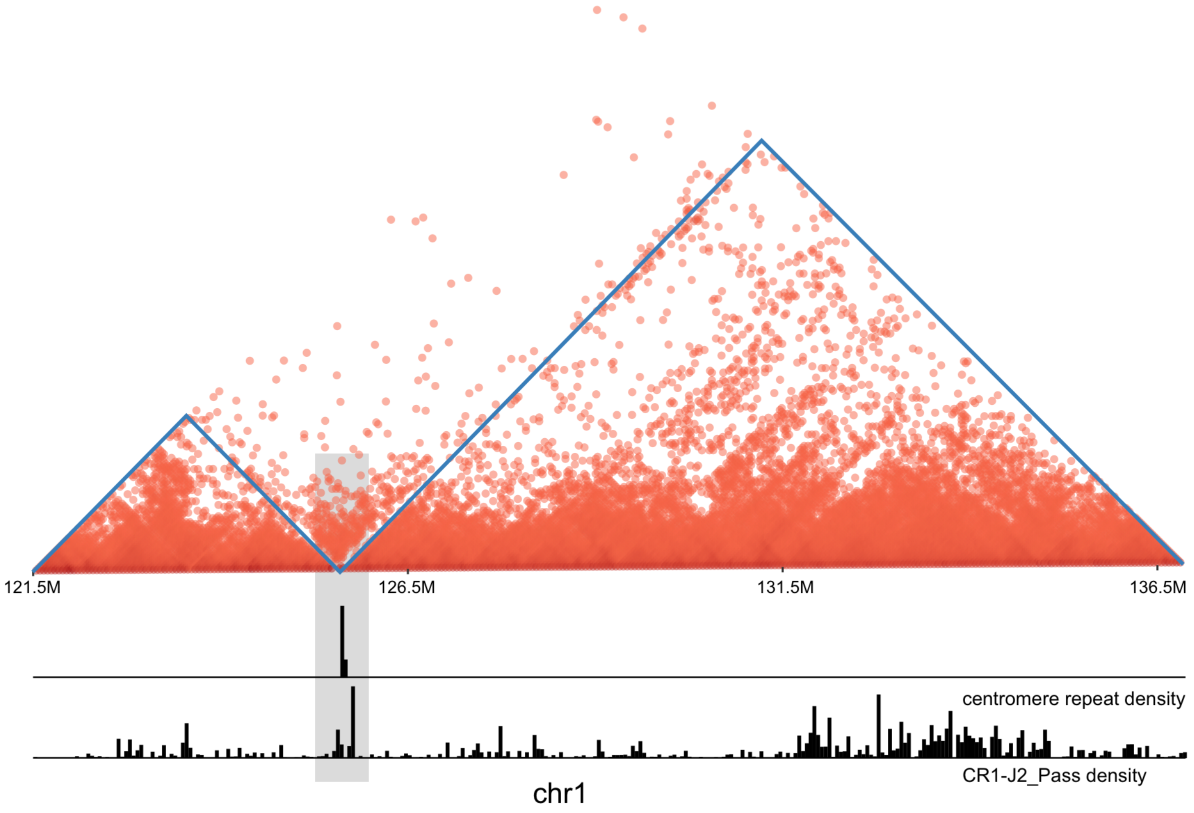


**Supplementary Figure S7 | An example centromere overlapped with the TAD boundary and enriched for CR1-J2_Pass repeats.** The Hi-C matrix plot on Chr1 was generated with ggplot2[1] packages with 50kb resolution. The CR1-J2_Pass density was calculated in 50kb non-overlapping windows. The region of the TAD boundary is shown in a light grey box.

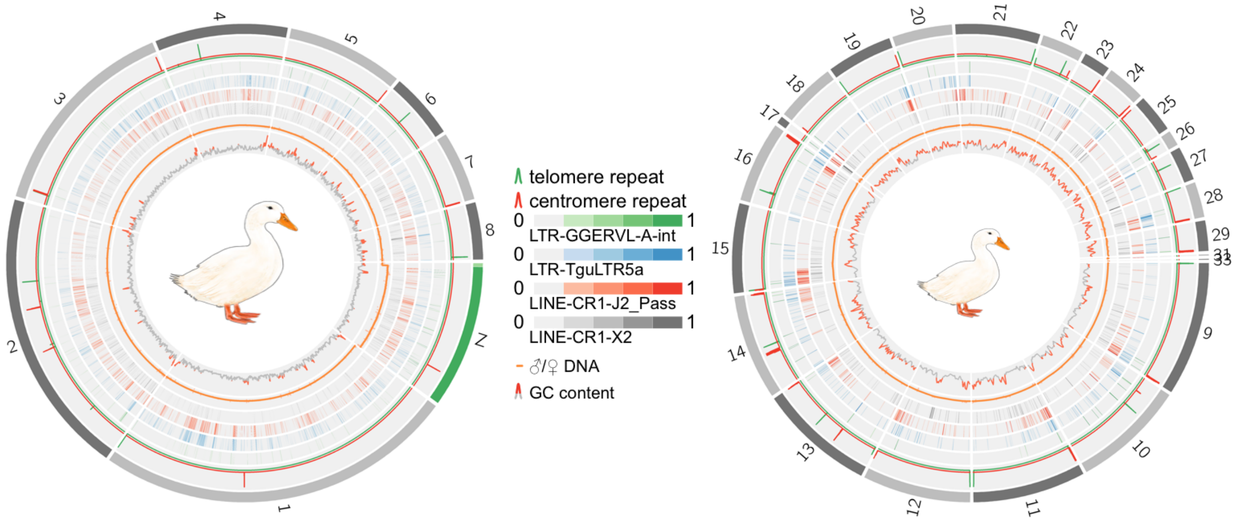


**Supplementary Figure S8 | Different repeats are enriched at centromeres of macro- and micro-chromosomes.** CR1-J2_Pass is enriched at the centromeres of both macro and micro-chromosomes. However, microchromosome centromeres are specifically enriched for the LTR repeat GGERVL-A-int. The LINE-CR1-X2 is also shown for comparison.

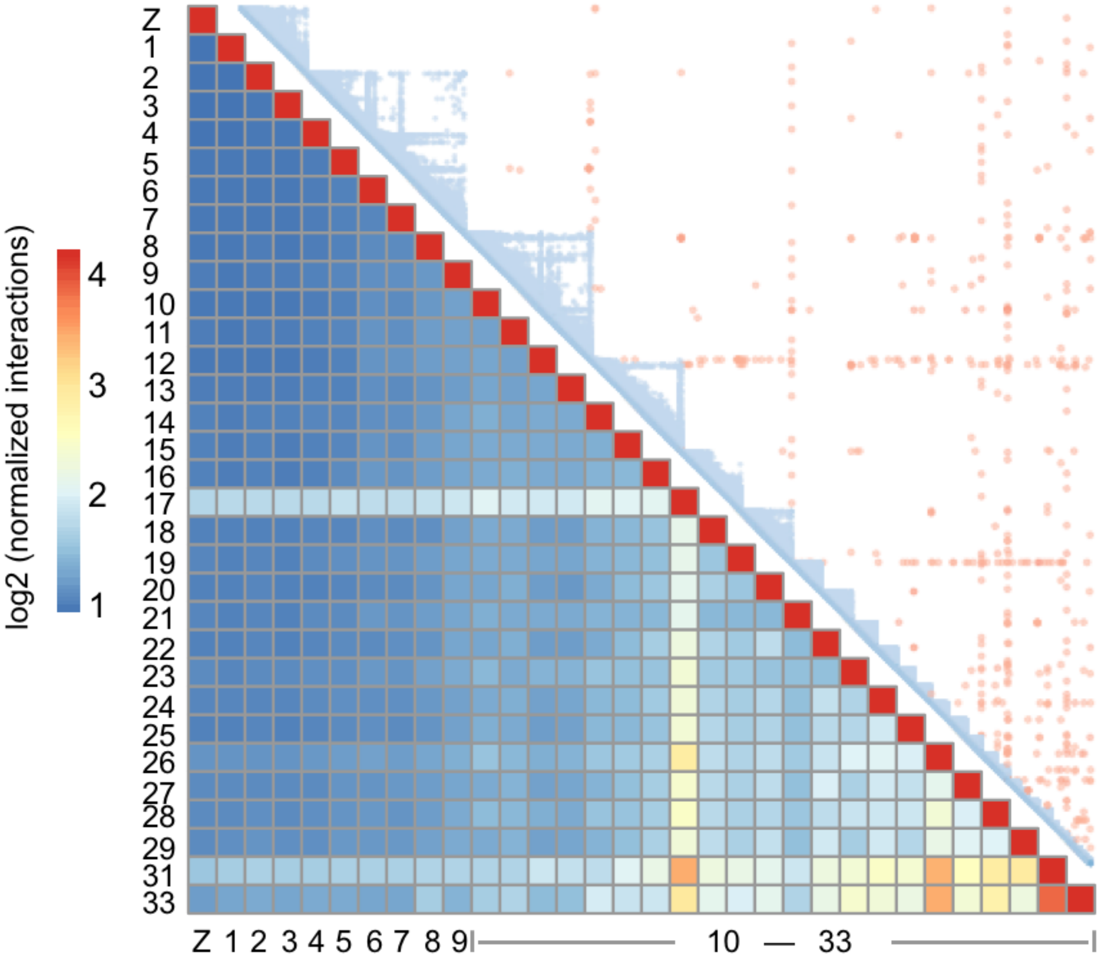


**Supplementary Figure S9 | Inter-chromosomal interactions.** Microchromosomes exhibit an unusual pattern of more frequent inter-chromosomal interactions measured by the Hi-C matrix (10kb resolution) compared to macrochromosomes. The blue triangles in the upper right indicate the intrachromosomal interactions, while the red dots indicate interchromosomal interactions. In the lower left heatmap, we showed the strength of such interactions measured by normalized Hi-C read pairs.

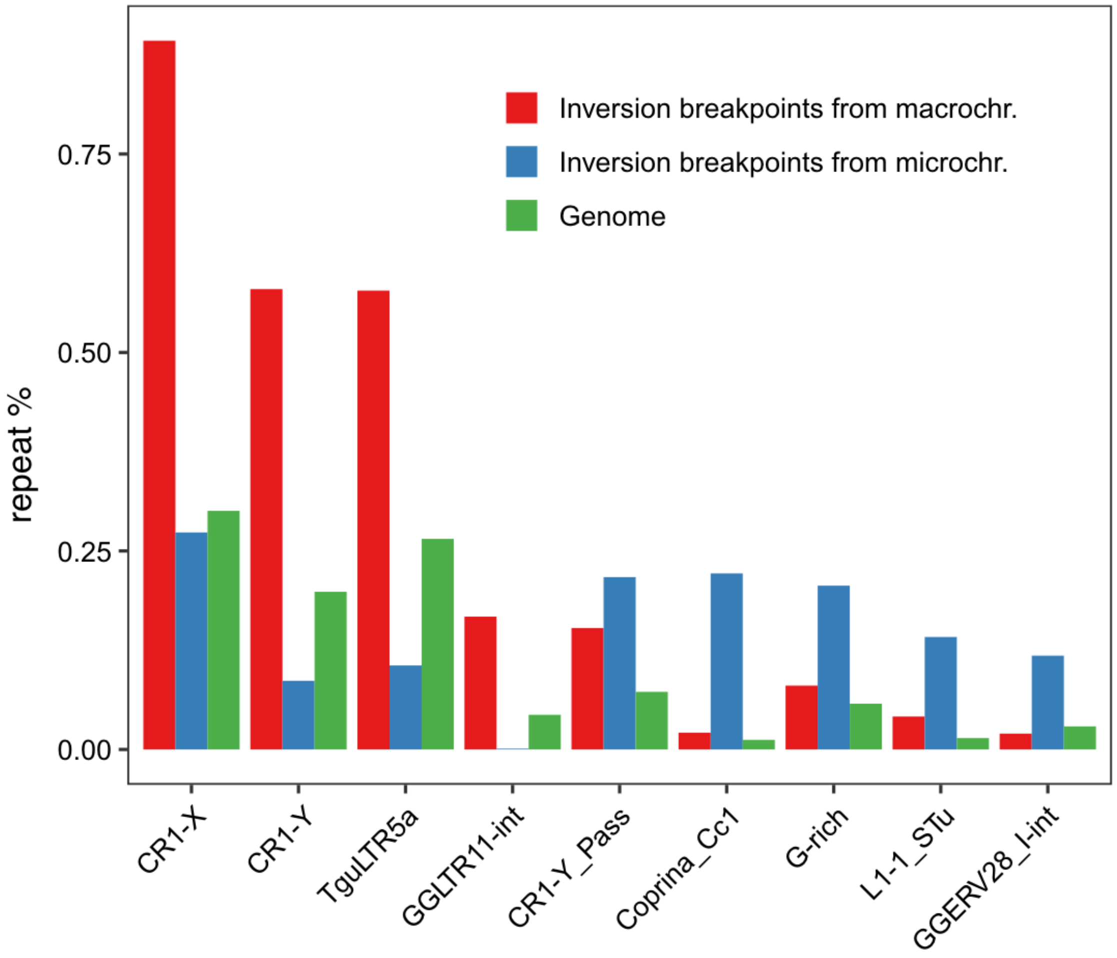


**Supplementary Figure S10 | Repeat enrichment at the inversion breakpoints.** For macro- vs micro-chromosomes, different repeats are enriched at their inversion breakpoints. Here we show the top 5 enriched repeats at the inversion breakpoints of macro- and micro-chromosomes. From left to right, the first five repeats are enriched at macrochromosomes and the last five repeats are enriched at microchromosomes. Within them, only CR1-Y_Pass is enriched for both macro- and micro-chromosomes.

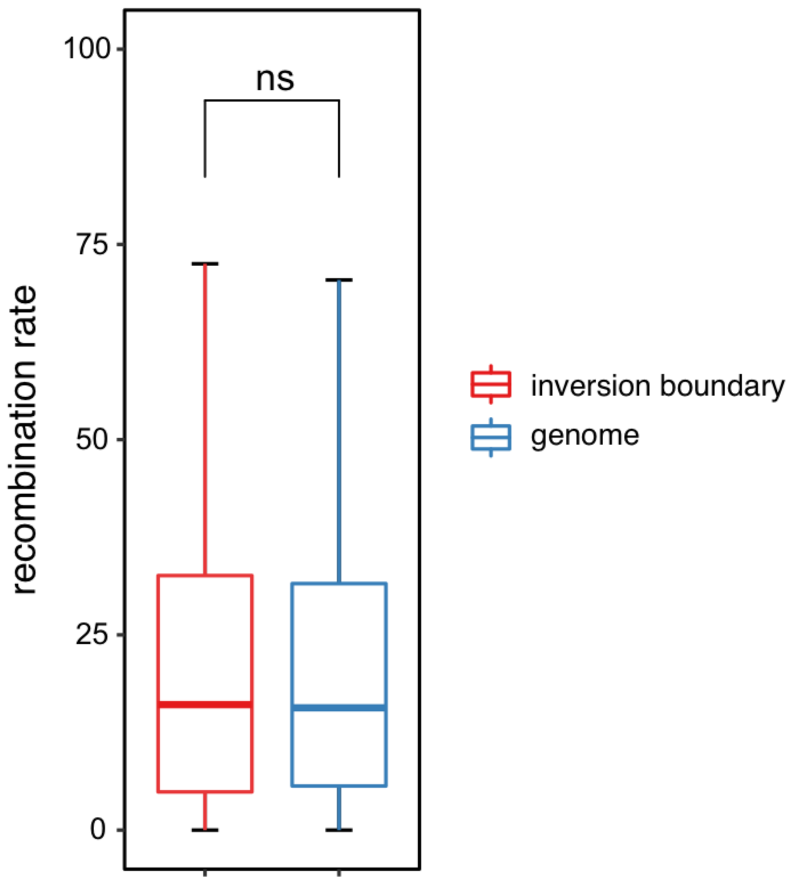


**Supplementary Figure S11 | Recombination rate at the inversion breakpoints.** The recombination rate between inversion breakpoints and the whole genome has no difference. “ns” indicates P = 0.533, measured by one-sided Wilcoxon signed rank test.

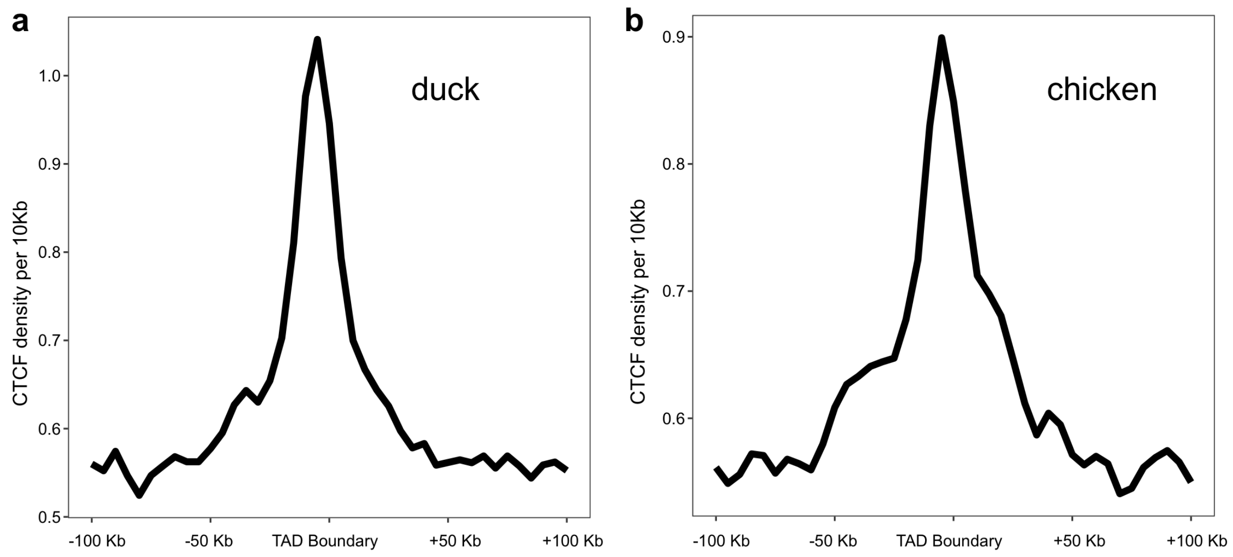


**Supplementary Figure S12 | CTCF is enriched at TAD boundaries in the duck and chicken genome.** CTCF density was calculated in 10kb non-overlapping windows along the up- and down-stream (100kb) regions of both chicken and duck TAD boundaries. For each window, the mean value of the CTCF density was used to generate the plots.

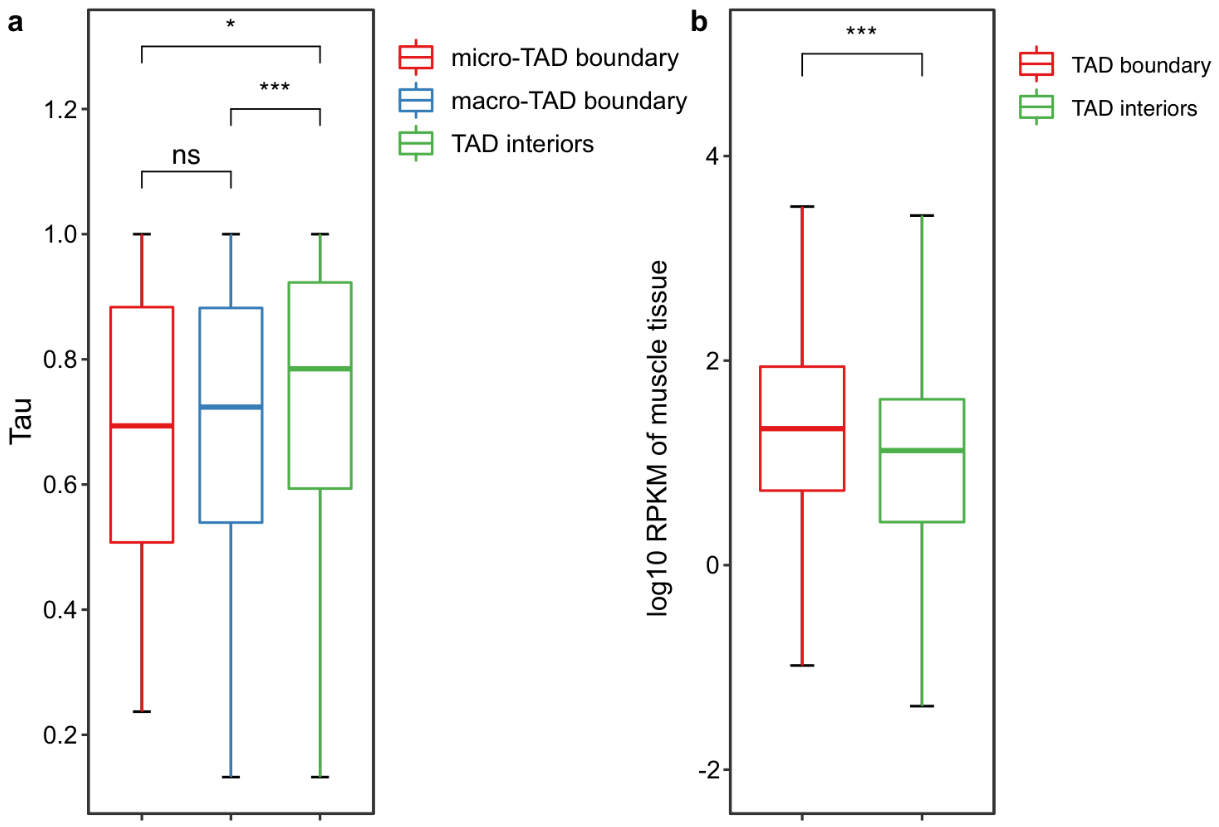


**Supplementary Figure S13 | TAD boundaries tend to be enriched for broadly expressed housekeeping genes in the Pekin duck. a.** The tau value (tissue specificity matrix) of genes located at TAD boundaries of both macro- and micro-chromosomes is significantly lower than those within TAD interiors, indicating a broader expression pattern of the genes, if any, located at the TAD boundaries. **b**. Expression levels of genes located at TAD boundaries are significantly higher than those within TAD interiors. "ns", P = 0.3299; . "*"; P = 0.02497;. "***",P < 2.2e-16; values determined by one-sided Wilcoxon signed rank test.

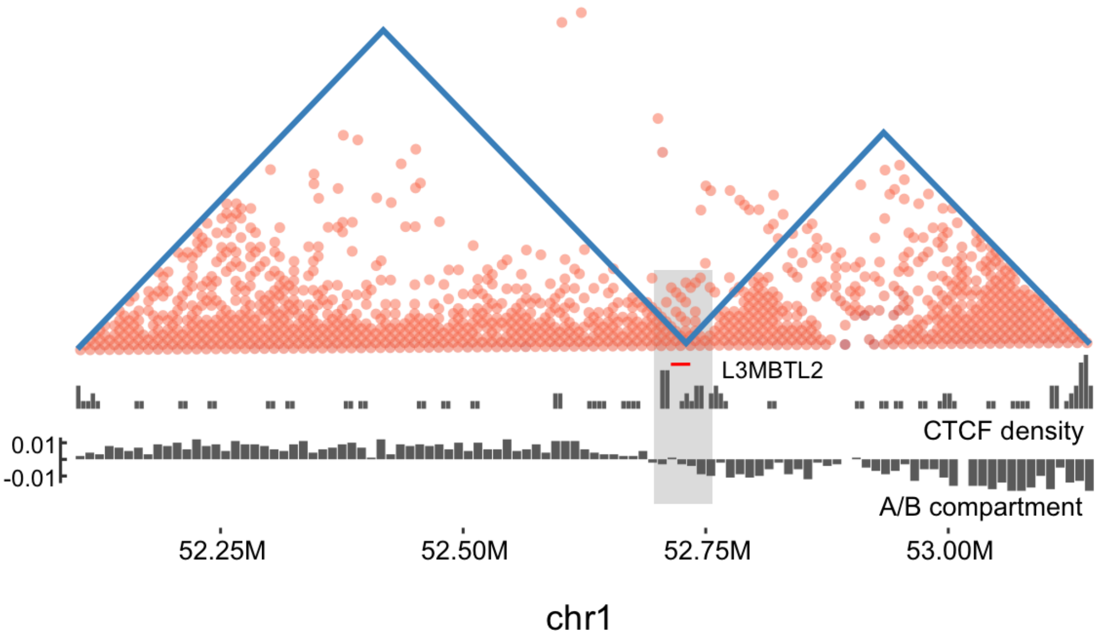


**Supplementary Figure S14 | An example of diverse types of TAD boundaries that overlap.** The Hi-C matrix plot is generated with ggplot2[1] packages with 50kb resolution. The CTCF density is calculated in 10kb non-overlapping windows. The A/B compartment is generated with a 10kb Hi-C matrix. The region of the TAD boundary is shown in a light grey box. This TAD boundary is enriched for putative CTCF binding sites, and also at the transition between the A/B compartments.

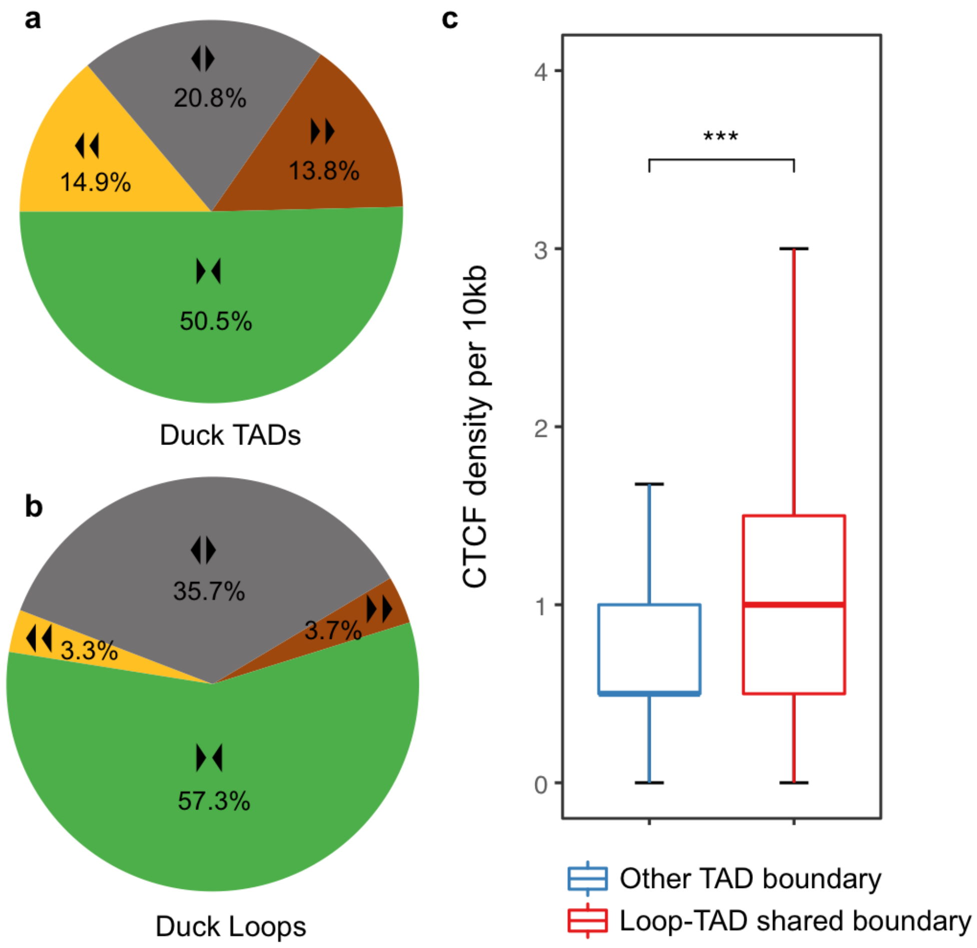


**Supplementary Figure S15 | DNA loop regions have a higher CTCF density and a higher percentage of paired CTCF sites in convergent orientation. a.** Pie chart shows four kinds of orientation (arrowheads and different colors, green color for the convergent CTCF site pairs) for the paired CTCF sites in the boundaries of Pekin duck TADs. **b.** Pie chart for the DNA loops. "***",P < 2.2e-16, measured by one-sided Wilcoxon signed rank test. **c.** The CTCF density of DNA loop boundaries is significantly higher than other TAD boundaries.

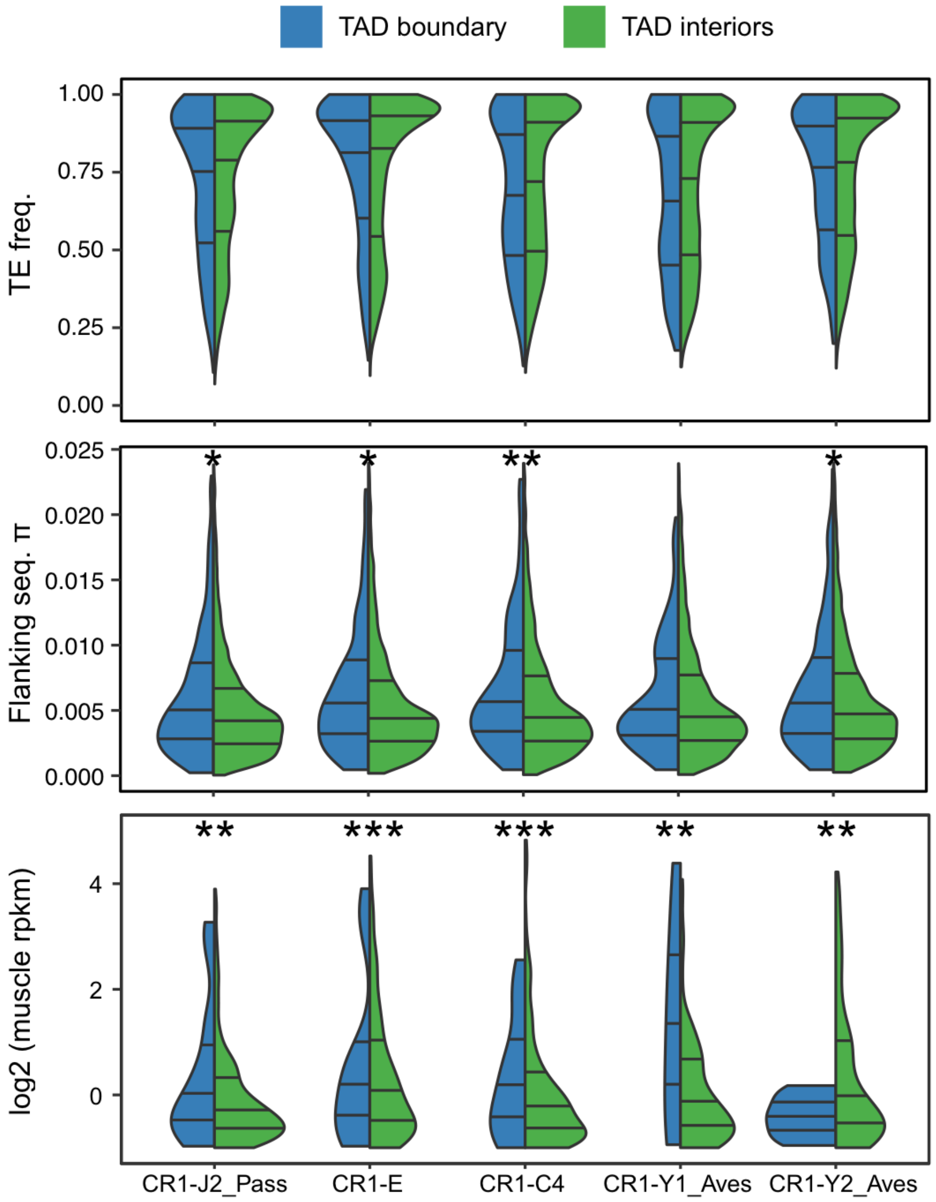


**Supplementary Figure S16 | TEs at the TAD boundaries.** Shown are the violin plots of the top 5 most enriched TEs at the TAD boundaries . They generally show a lower population frequency (top), a higher level of segregating sequence polymorphism in their flanking sequences (middle), and a higher expression level in the muscle tissue (bottom) compared to the same TEs in TAD interiors (blue vs. green). *,P < 0.05; **,P < 0.005,;***, P < 0.0005. One-sided Wilcoxon signed rank test.

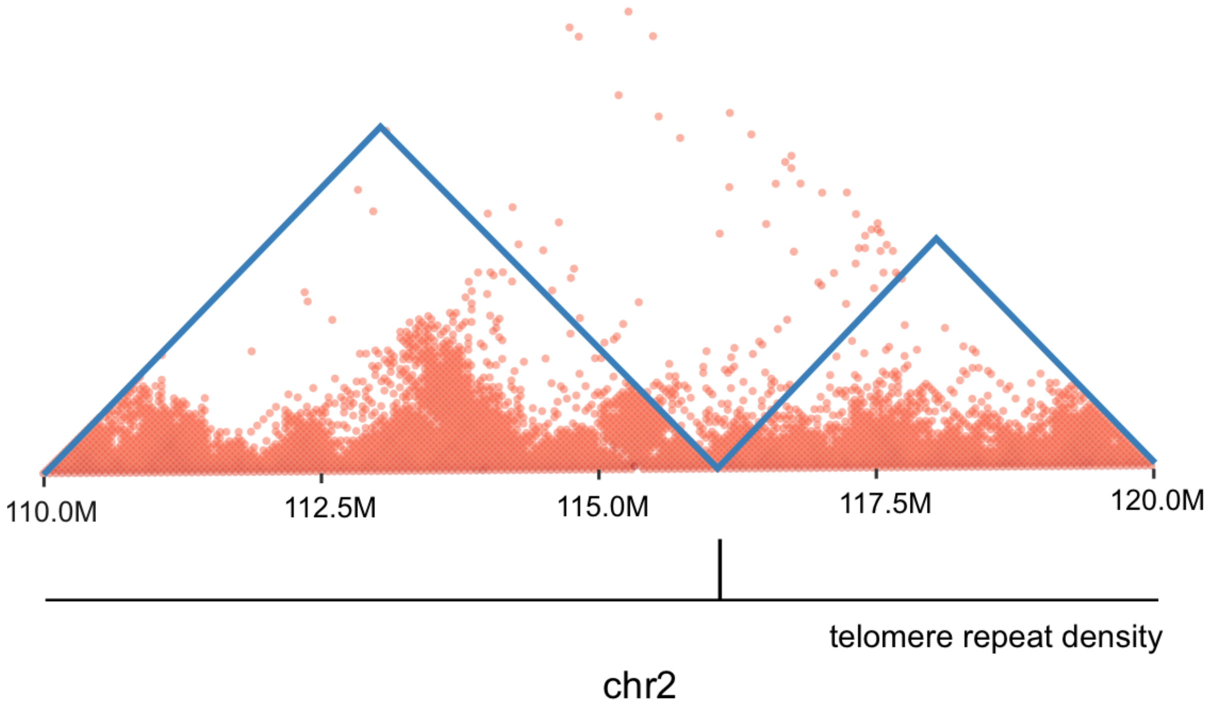


**Supplementary Figure S17 | An example of interstitial telomere sequences that are overlapped with a TAD boundary.**

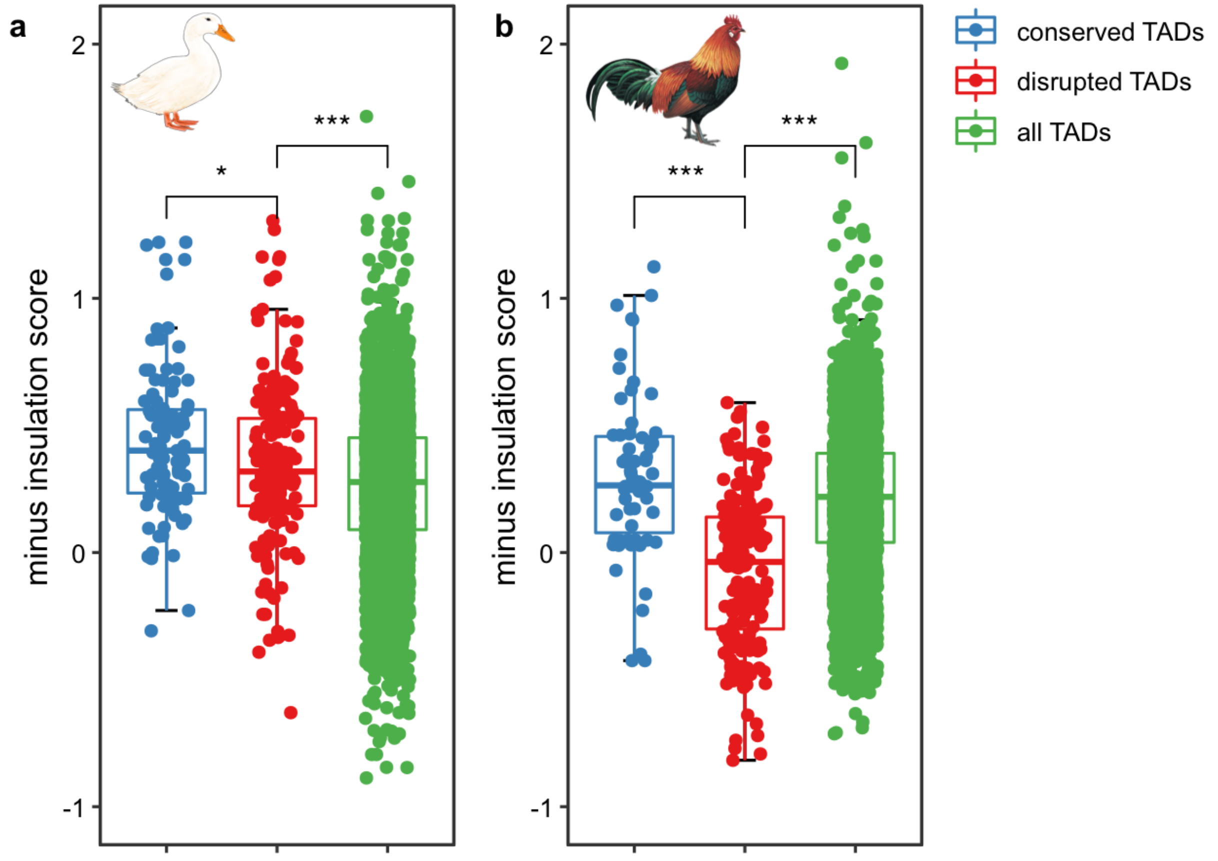


**Supplementary Figure S18 | Disrupted TADs tend to have a higher insulation score. a.** TAD insulation scores in duck. **b.** TAD insulation scores in chicken. Conserved TADs mean the TAD boundaries were shared between duck and chicken. Disrupted TADs refer to the TADs that were influenced by the duck- or chicken-specific inversions. All TADs include the conserved, disrupted, and others. Note the negative scores of insulation scores were shown. *, P < 0.05; **,P < 0.005; *** ,P < 0.0005. Values measured by one-sided Wilcoxon signed rank test.

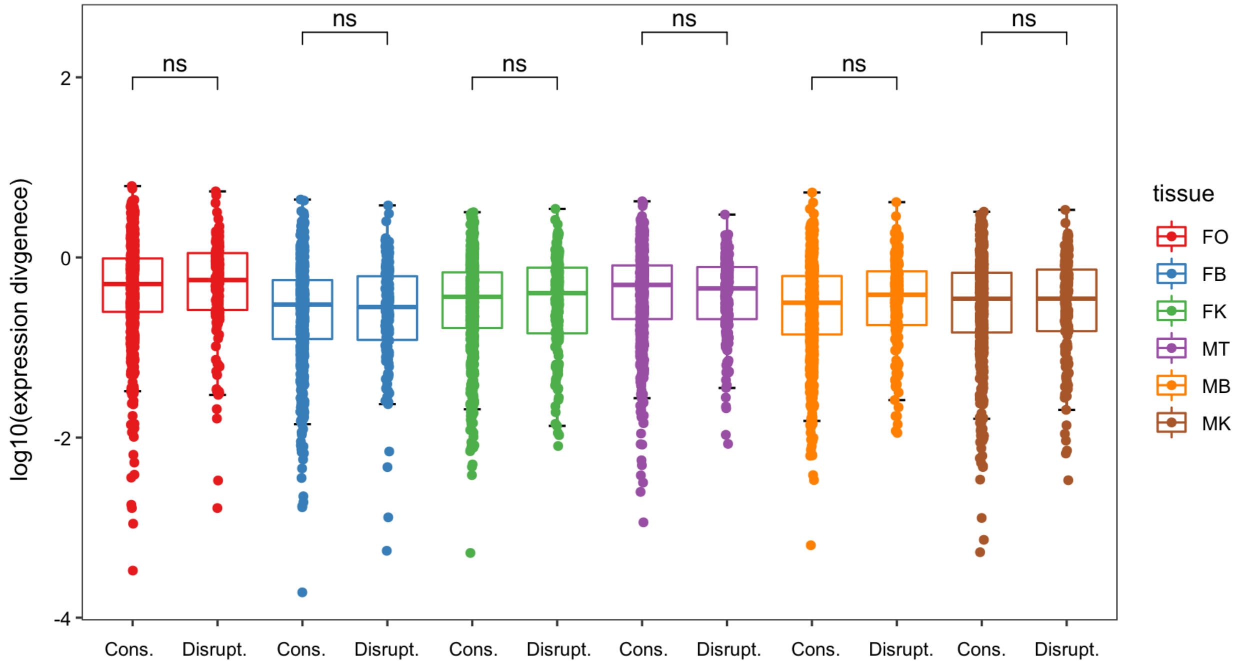


**Supplementary Figure S19 | No significant difference of expression divergence between disrupted and conserved TAD domains.** Disrupted TAD domains mean the TADs were created by the duck- or chicken-specific inversions. Conserved TADs mean the TADs were shared between duck and chicken. "ns" indicates $P > 0.05$, measured by one-sided Wilcoxon signed rank test. FO=Female Ovary, FB=Female Brain, FK=Female Kidney, MT=Male Testis, MB=Male Brain, MK=Male Kidney.

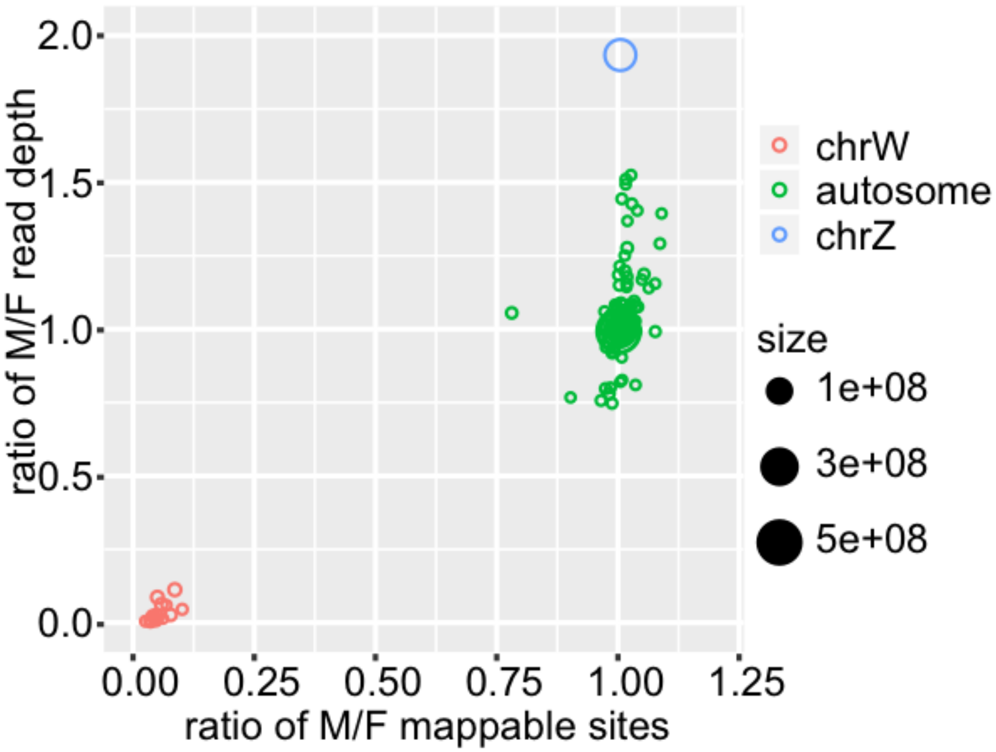


**Supplementary Figure S20 | Sex chromosomes show a different coverage pattern between sexes.** Z and W chromosomes were validated by calculating the male vs. female (M/F) ratios of read depth and mappable sites. The chrW scaffolds have a ratio of M/F read depth, and that of M/F mappable sites close to 0. The chrZ has a ratio of M/F read depth close to 2, and a M/F mappable sites ratio close to 1.

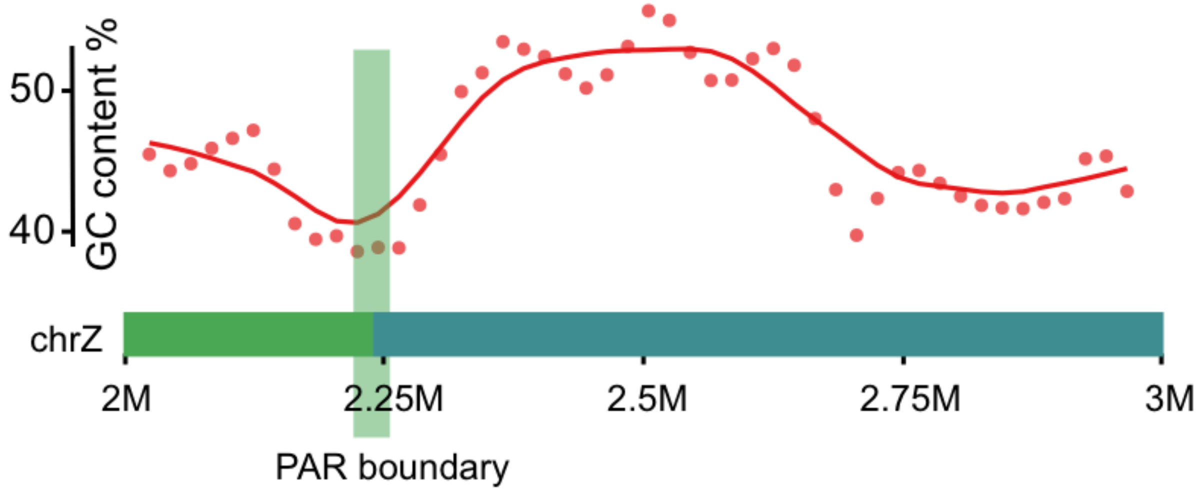


**Supplementary Figure S21 | GC shift along PAR boundary.** The GC content was calculated in 50kb windows with 25kb overlapped along the Z chromosome of Pekin duck. The highlighted region at 2.25 Mb is the boundary between PAR and S3 region of duck on the chrZ.

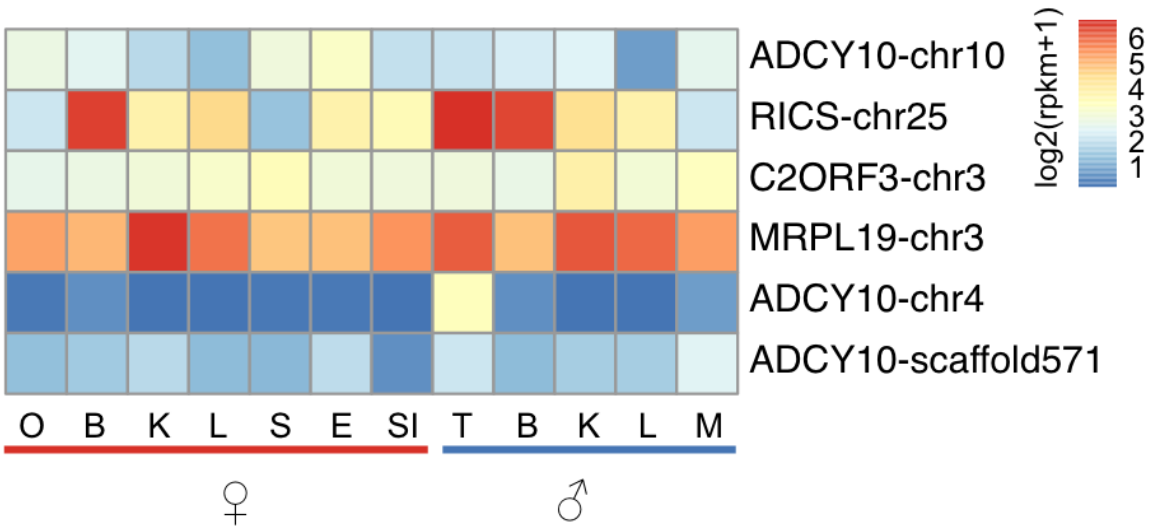


**Supplementary Figure S22 | Chicken Z-amplicon orthologs' expression in Pekin duck.** We have not found the similar tandem arrays of testis genes on the Z chromosome of chicken in duck. RPKM are normalized RNA-seq read expression levels, with red (6) being higher. These testis specific genes are on the autosomes indicated in the x-axis. O=Ovary, B=Brain, K=Kidney, L=Liver, S=Spleen, E=Egg shell gland, SI=Small intestine, T=Testis, M=Muscle.

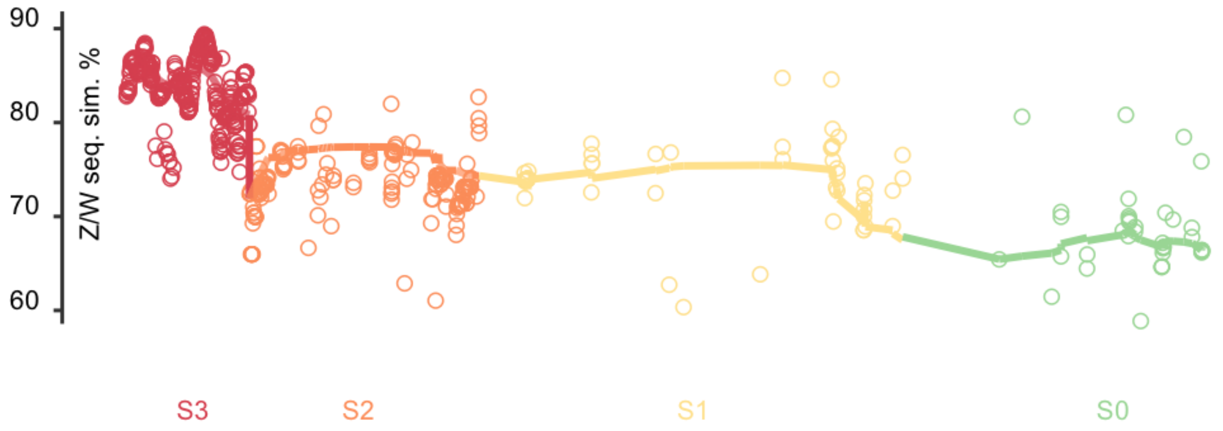


**Supplementary Figure S23 | Sequence similarity between duck W and emu Z.** The duck W scaffolds were mapped to emu Z chromosomes. Each dot represents one 50kb long non-overlapping window. Within each stratum, chrW scaffolds of similar levels of sequence divergence are clustered and separated from the neighbouring strata with a different divergence level. The strata identification is detailed in the "Evolutionary strata" section of Methods part.

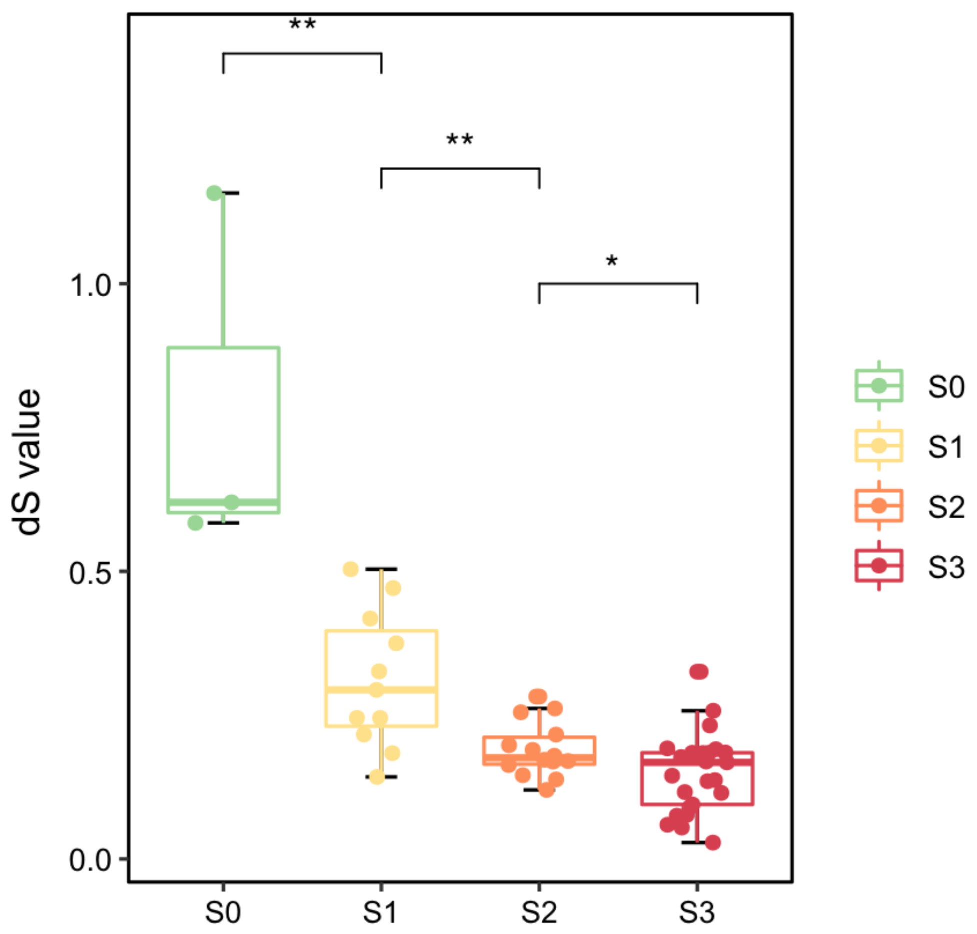


**Supplementary Figure S24 | The distribution of pairwise dS values of duck sex chromosomes.** The genes within each stratum show a consistent gradient of synonymous substitution rates, from old (S0) to young (S3) stratum. *,P < 0.05; **, P < 0.005; measured by one-sided Wilcoxon signed rank test.

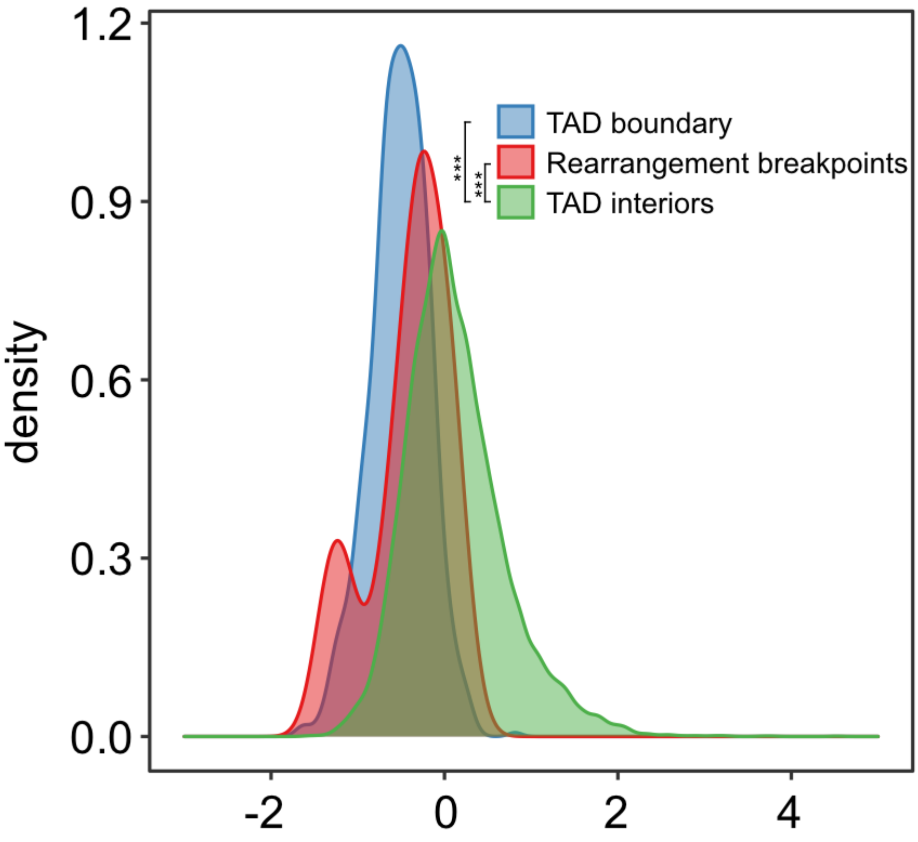


**Supplementary Figure S25 | Rearrangement breakpoints between duck and emu chrZ tend to have a low insulation score.** The x axis is the insulation score. ***, P < 2.2e-16, measured by one-sided Wilcoxon signed rank test.

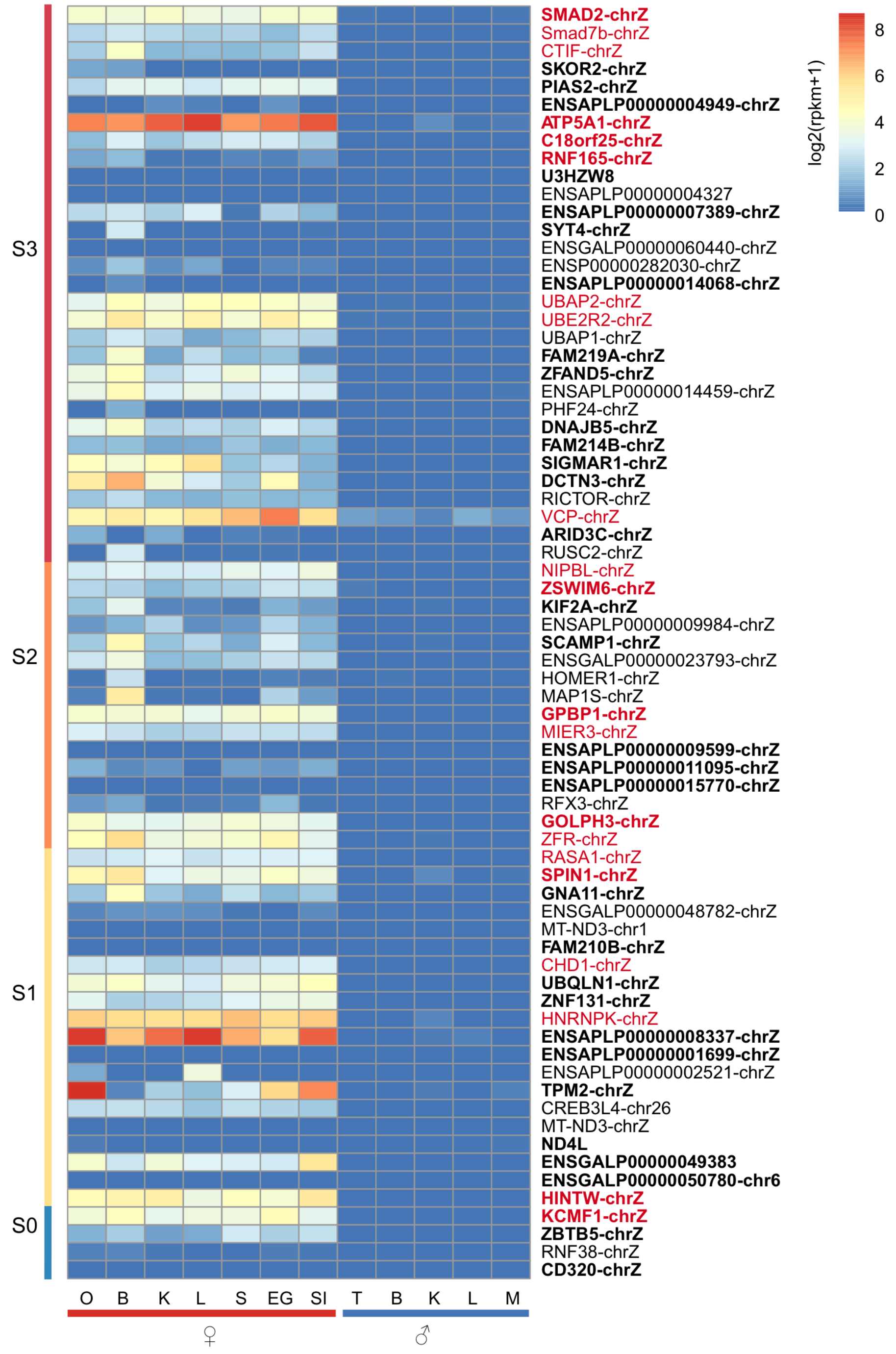


**Supplementary Figure S26 | Expression pattern of duck chrW genes.** Genes with intact ORFs are shown in bold text. Genes without intact ORFs are shown in regular text. Genes shared with chicken chrW are shown in red color. RPKM are RNASeq read expression levels, with red being higher. Note, lack of expression in males, as expected. O=Ovary, B=Brain, K=Kidney, L=Liver, S=Spleen, E=Egg Shell ganld, SI=Small intestine, T=Testis, M=Muscle.

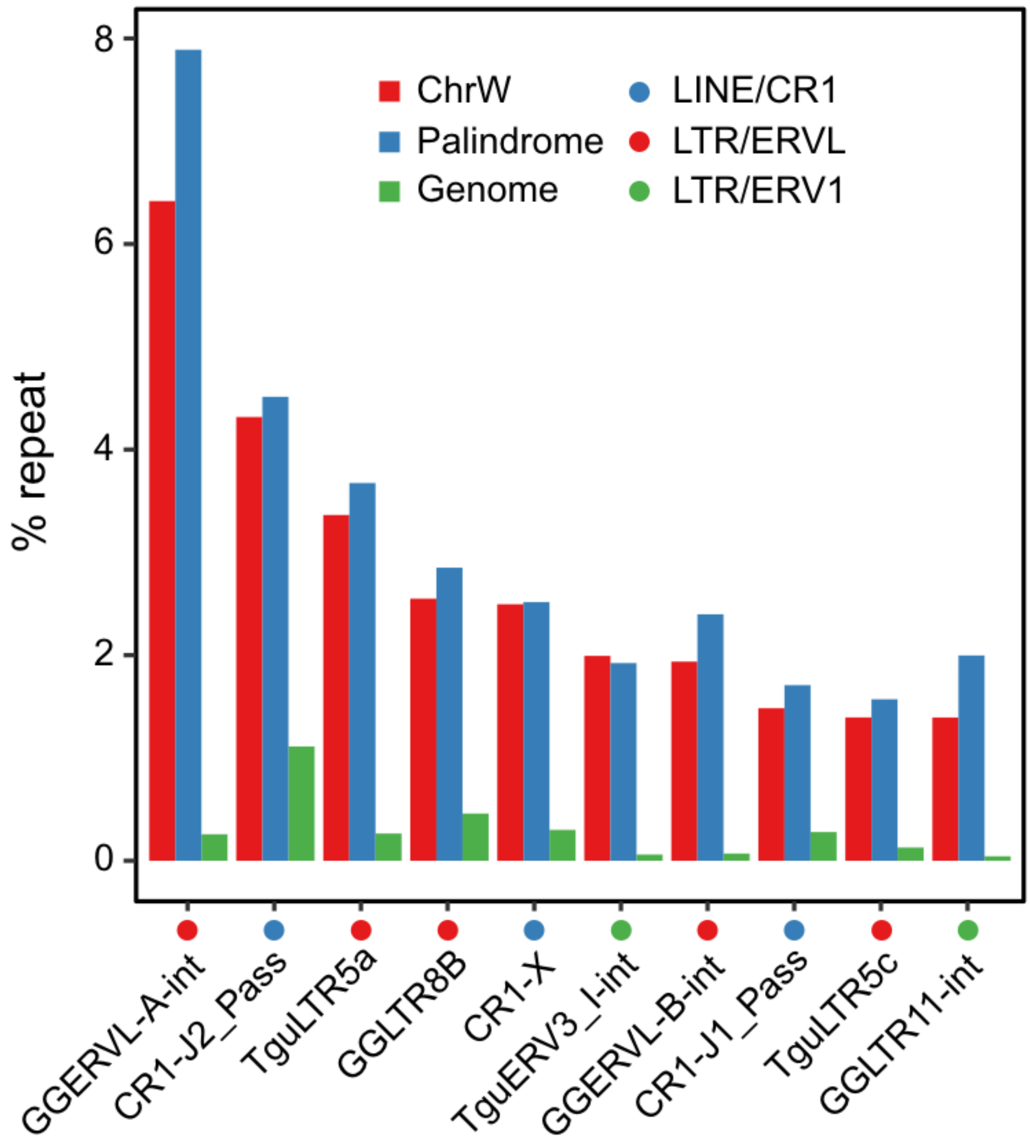


**Supplementary Figure 27 | Repeats enriched in duck chrW.** Shown are the top 10 most abundant repeats on the chrW of Pekin duck.

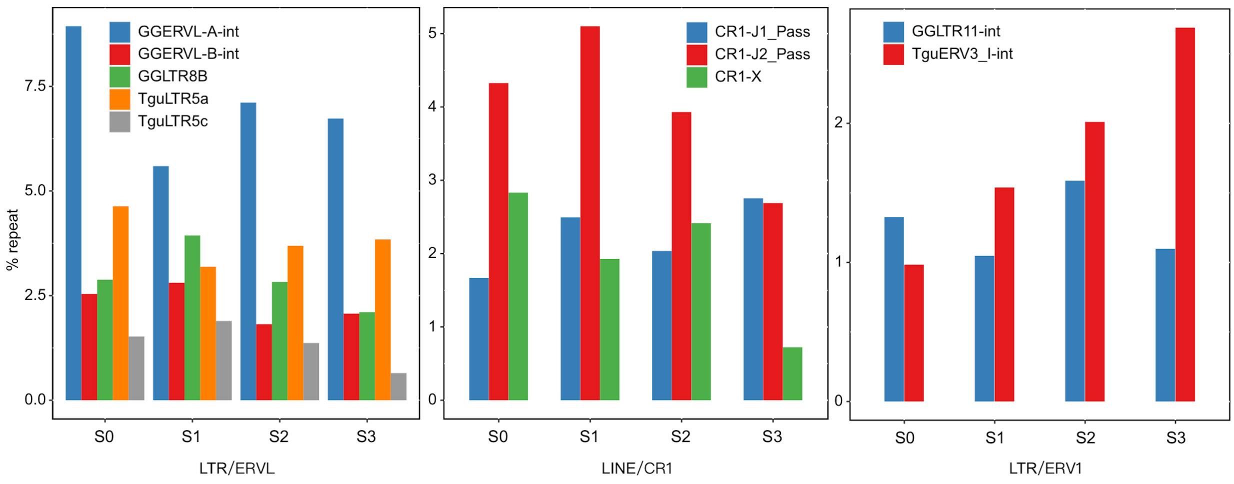


**Supplementary Figure S28 | Repeats enriched at different evolutionary strata.** Different TE families exhibit opposing trends of colonizing the different age of evolutionary strata. TE families that have been propagating since the ancestor of Neoaves (e.g., CR1-J2_Pass) are more enriched in the older strata, while TE families that were specifically propagated in the duck (e.g., TguERV3_I-int) are more enriched in the younger strata.
